# The composition of the perinatal intestinal microbiota in horse

**DOI:** 10.1101/726109

**Authors:** A Husso, J Jalanka, MJ Alipour, P Huhti, M Kareskoski, T Pessa-Morikawa, A Iivanainen, M Niku

**Author notes:** Antti Iivanainen and Mikael Niku jointly supervised this work. Correspondence to: MN and AI.

## Abstract

The establishment of the intestinal microbiota is critical for the digestive and immune systems. We studied the early development of the microbiota in horse, a hindgut fermenter, from birth until 7 days of age, by qPCR and 16S rRNA gene amplicon sequencing. To evaluate initial sources of foal microbiota, we characterized dam fecal, vaginal and oral microbiotas. We utilised an amplicon sequence variant (ASV) based pipeline to maximize resolution and reproducibility. Stringent ASV filtering based on prevalence and abundance in samples and controls purged reagent contaminants while preserving intestinal taxa. The newborn rectum contained small amounts of diverse bacterial DNA, with a profile closer to mare feces and vagina than mouth. 24 hours after birth, the intestine was colonized by Firmicutes and Proteobacteria, some foals dominated by a single genus. At day 7, the phylum-level composition resembled adult feces but genera were different. The mare vaginal microbiota contributed to 24 h and 7 day microbiotas. It contained few lactobacilli, with *Corynebacterium*, *Porphyromonas*, *Campylobacter* and *Helcococcus* as the most abundant genera. In the oral mucosa, *Gemella* was extremely abundant. Our observations suggest that bacteria or bacterial components translocate to the equine fetus, but the intestinal microbiota changes rapidly after birth.

## Introduction

Intestinal microbiota is required for the development and function of the digestive and immune systems. The first microbes are obtained from the mother early in life^1–3^. The maternal microbiota affects the development of the intestinal immune system already in the fetal period^4^. This is mediated by circulating microbial metabolites^4^ and possibly also by small numbers of microbial cells translocating to the fetus via placenta^5–7^. The major microbial colonization occurs at birth. Postnatally, the intestinal microbiota goes through a rapid development, with changing bacterial composition, diversity and abundances, before reaching adulthood where the microbiota is relatively stable^8–10^. Early exposure to a diverse microbiota promotes the maturation of an immune system which effectively protects from infection while tolerating commensal microbes^11–13^.

Gut microbiota is especially important to herbivores, as they rely on bacterial fermentation to digest plant fibers to volatile fatty acids, which are their major energy source. In horse (*Equus ferus caballus)*, the microbial fermentation occurs in the enlarged hindgut: colon and caecum^14,15^. These compartments harbor extremely abundant and diverse microbiotas, the compositions of which are critical to the equine health^16^. Horses are susceptible to gastrointestinal complications induced by dietary changes, antimicrobials and infections, such as colic and colitis^15,17,18^.

The foal’s first week of life post-partum is considered a critical period, with increased morbidity and mortality in respiratory diseases, enteritis and sepsis^19–21^. Mild noninfectious “foal heat diarrhea” is common in neonatal foals during the first weeks^19^. It is probably associated with the ongoing microbial colonization of the gut and hypersecretion of the small intestinal mucosa. More serious enteritis and enterocolitis is caused by bacterial or viral infections or mucosal injury^19^. Septicaemia is typically associated with a failure of the transfer of colostral immunoglobulins^22^. Understanding the origins and development of the equine intestinal microbiota at early age may offer insight into the mechanisms of gastrointestinal diseases and offer new solutions for prevention of foal morbidity and mortality^19^. The development of the equine intestinal microbiota has been previously investigated by sequencing^23–25^. However, these studies did not include meconium samples collected immediately at birth, or did not utilise negative controls recommended in the analysis of low-biomass microbiota samples^26^.

In this study, we aimed to investigate the early development of the foal intestinal microbiota over the first postnatal week, starting at birth. We compared the foal microbiota to the fecal, vaginal and oral microbiotas of their dams. These locations have been regarded as potential initial sources of the early microbiota^9,27,28^. We used 16S rRNA gene amplicon sequencing and quantitative PCR to characterize the composition and abundance of the various microbiotas. We utilised a bioinformatics pipeline which has been shown to improve the resolution and reproducibility of amplicon sequencing^29,30^. This enabled stringent data filtering to efficiently remove potentially contaminating sequences, based on their prevalence and relative abundance in samples and several types of negative controls.

## Results

### Quantification of 16S rRNA gene copy numbers in foals and negative controls

To estimate the bacterial loads in the foals and negative controls, we analysed the 16S rRNA gene copy numbers using qPCR. In newborn foals, there were on average 7.8 × 10^4^ copies (SD = 1.2 × 10^5^) of 16S rRNA genes per sampling swab (Fig. 1). This was significantly more (P = 0.006) than in the field controls (sampling instrument exposed to farm air + extraction kit; 4.3 × 10^3^ copies per swab; SD = 7.6 × 10^2^). Instrument controls (sampling instrument + extraction kit) contained 3.6 × 10^3^ copies per swab (SD = 7.2 × 10^2^) and no-template PCR controls 9.7 × 10^2^ copies per sample (SD = 3.4 × 10^2^).

**FIGURE 1.**
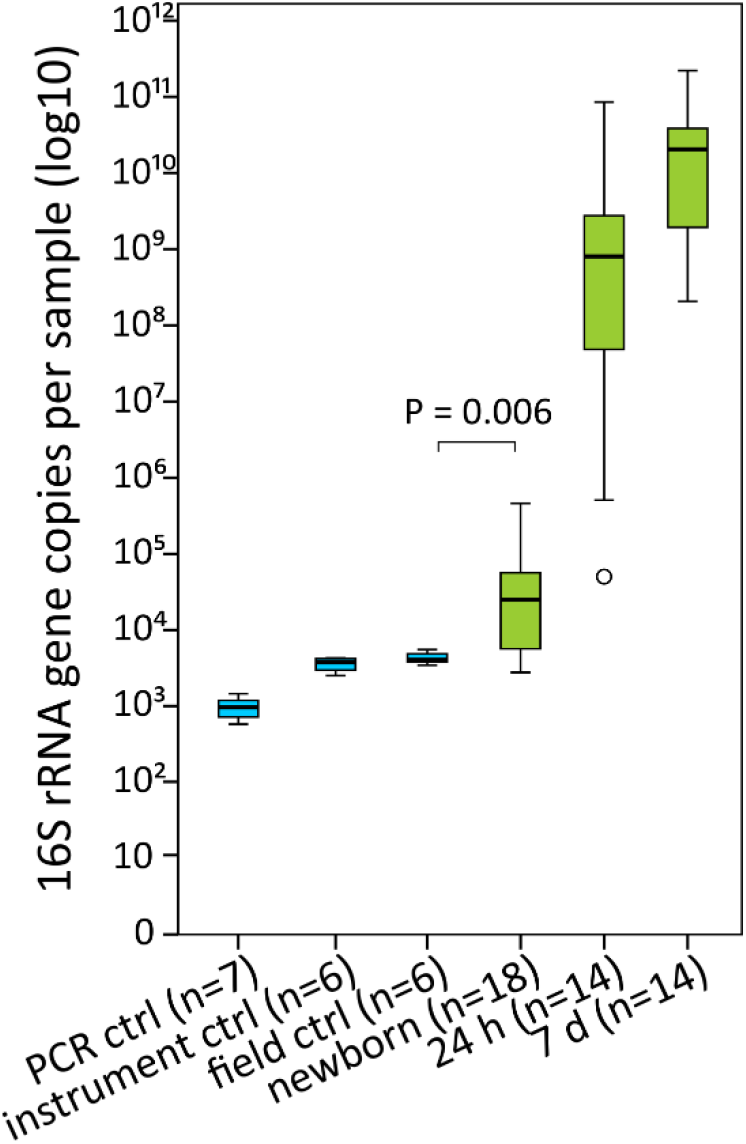
16S rRNA gene copy numbers per sample. Blue colour indicates the negative controls and green the foal rectal samples. The boxes represent the interquartile ranges (IQR) containing 50% of samples. The horizontal line in a box indicates the median. Whiskers show maxima and minima within 1.5 × IQR. The circle indicates an outlier.

Already 24 h after birth, there was a four-log increase in the bacterial load (7.4 × 10^9^ copies per swab, SD = 2.3 × 10^10^). There was a further increase at the sample collected 7 days after birth, to a mean of 4.2 × 10^10^ copies (SD = 6.2 × 10^10^; Fig. 1).

### Sequencing controls and decontamination of 16S rRNA amplicon sequencing data

We used a commercial microbial community composition standard to control the entire analytical pipeline from DNA extraction to bioinformatics. The observed composition and abundances matched the expected composition provided by the manufacturer (Supplementary Fig. 1). *Staphylococcus aureus*, *Listeria monocytogenes* and *Lactobacillus fermentum* were correctly classified to species level, with an exception of 16% *Staphylococcus* reads classified to the genus level. The other bacteria were correctly classified to the genus level.

We utilised several types of negative controls in the 16S rRNA gene sequencing to minimize the risk of false positive observations: PCR controls, DNA extraction controls, instrument controls and field controls^26^. Stringent filtering of the 16S rRNA gene sequencing data was performed to remove amplicon sequence variants (ASVs) potentially originating from contaminants. The filtering was based on comparison of the prevalence and relative abundance of each ASV in samples and negative controls, as described in the Methods section. On average, the decontamination procedure removed 99.9% (SD = 0.186) of sequence reads from the negative controls, 84.0% (SD = 24.3) from newborn rectal samples, 10.2% (SD = 27.1) from 24 h samples, 4.36% (SD = 3.09) from 7 d samples and 1.98% (SD = 6.50) from the various dam samples (Fig. 2). Most of the removed ASVs were classified as *Ralstonia*, a common reagent contaminant^26^. Across all animal samples, 99.8% of *Ralstonia* reads were removed. In contrast, for ASVs classified as typical intestinal genera, which are less likely reagent contaminants, only 0.179% of reads were removed across all samples, and 3.89% in the newborn samples (for details, see Supplementary Materials).

**FIGURE 2.**
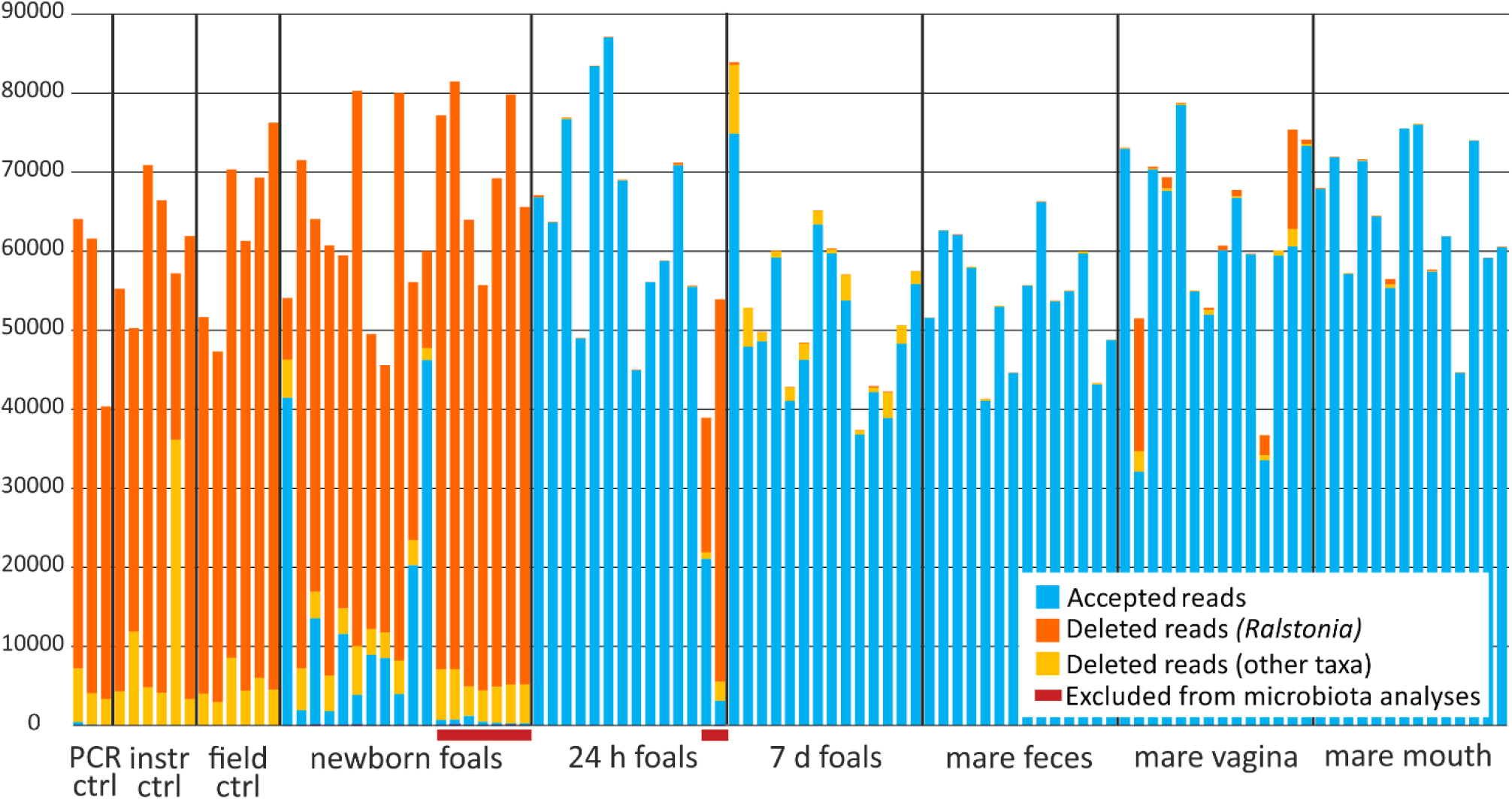
Accepted and rejected 16S rRNA gene sequence reads and 16S rRNA gene copy numbers per sample. Accepted reads are indicated as blue. Deleted reads are indicated as yellow-orange, with reads classified as Ralstonia in orange. Seven newborn samples and two 24 h samples were excluded from further analysis due to low quality (red bars). Negative control data processed with the newborn data is shown.

After the data decontamination, seven newborn samples were excluded from further microbiota composition analyses due to small number of accepted reads (< 1500; Fig. 2). In two of these, the total DNA concentrations were below Qubit detection limit. This suggests inadequate sampling, as in most cases the samples contain measurable host DNA from the intestinal mucosa. Two 24 h samples were also excluded due to low quality (small number of accepted reads and unusual microbiota composition). Also in these cases, the total DNA concentration was low or undetectable.

An overview of the raw and decontaminated data is shown in Table 1. All further analyses were performed using the decontaminated data.

**TABLE 1.** Overview of the 16S rRNA gene sequencing data before and after decontamination. Mean values (and standard deviations) for read and ASV counts in raw and decontaminated data are shown. Negative control data processed with the newborn data is shown.

|  | Newborn rectum | 24 h feces | 7 d feces | Mare feces | Mare mouth | Mare vagina | PCR control | Instrument control | Field control |
| --- | --- | --- | --- | --- | --- | --- | --- | --- | --- |
| **RAW DATA** |  |  |  |  |  |  |  |  |  |
| *N* | 18 | 14 | 14 | 14 | 14 | 14 | 3 | 6 | 6 |
| Raw reads | 66665<br>(9921) | 62657<br>(13583) | 53740<br>(11447) | 54666<br>(7346) | 64303<br>(8736) | 63612<br>(11207) | 55363<br>(10639) | 60373<br>(6927) | 62733<br>(10381) |
| Raw ASVs | 154<br>(143) | 82<br>(56) | 163<br>(41) | 1360<br>(146) | 79<br>(51) | 358<br>(392) | 46<br>(15) | 49<br>(61) | 73<br>(13) |
| **DECONTAMINATED DATA** |  |  |  |  |  |  |  |  |  |
| *N* | 11 | 12 | 14 | 14 | 14 | 14 | 3 | 6 | 6 |
| Decontaminated reads | 10645<br>(11605) | 65149<br>(12533) | 51212<br>(10231) | 53936<br>(7302) | 64089<br>(8816) | 60131<br>(13273) | 213<br>(218) | 12<br>(22) | 5<br>(0.5) |
| Decontaminated ASVs | 162<br>(117) | 61<br>(29) | 142<br>(34) | 1235<br>(132) | 66<br>(41) | 300<br>(345) | 3<br>(1.5) | 0.3<br>(0.5) | 0.3<br>(0.5) |

### Characterization of the newborn foal rectal microbiota

The newborn rectal microbiota consisted primarily of Proteobacteria, Firmicutes, Actinobacteria and Bacteroidetes (Fig. 3 and Table 2). The most prevalent bacterial genera in newborns are shown in Supplementary Table 1. These included typical animal-associated taxa such as *Staphylococcus, Lactobacillus, Bacillus, Streptococcus, Corynebacterium* and *Sphingomonas*. *Staphylococcus* was very abundant in some of the foals (up to 39% of all reads). The representative sequences of the most common staphylococcal, streptococcal and *Lactobacillus* ASVs were 100% identical to equine-associated species (*Staphylococcus equorum, Streptococcus equinus, Lactobacillus equi* and *Lactobacillus equigenerosi*). Some of the genera observed in newborns, such as *Ralstonia* and *Methylobacterium*, may represent remaining reagent contaminants.

**FIGURE 3.**
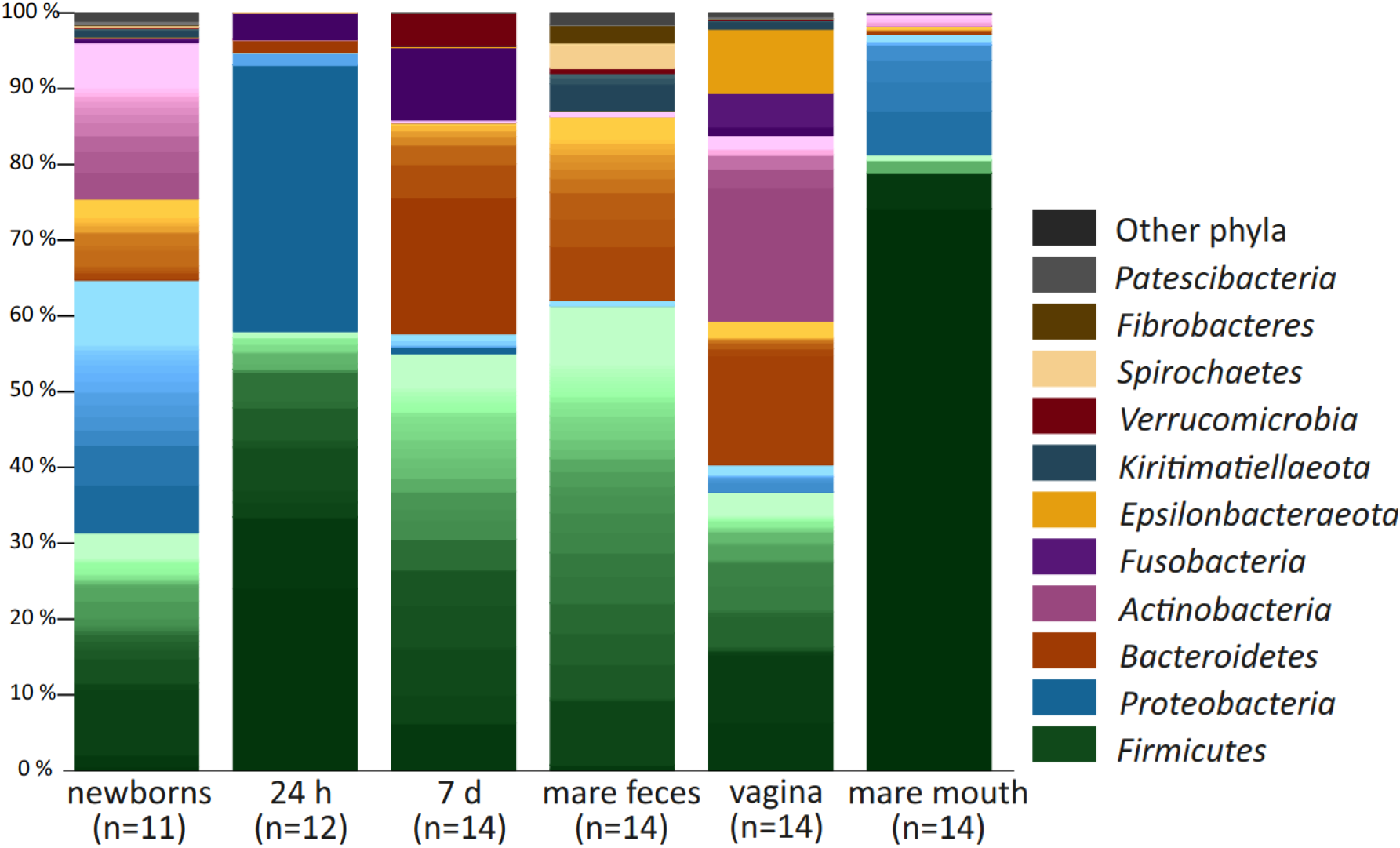
Average microbiota compositions of foal rectal samples and mare fecal, vaginal and oral samples. Main colours indicate bacterial phyla. Within phyla, shades indicate individual genera. The lightest shades of each phylum show the combined abundance of the least abundant genera (< 0.5% of total).

**TABLE 2.** Relative abundances of the major bacterial phyla and Shannon diversity indices in foal rectal samples and mare fecal, vaginal and oral samples (mean values and standard deviations).

|  | Newborn | 24 h | 7 d | Mare feces | Mare vagina | Mare mouth |
| --- | --- | --- | --- | --- | --- | --- |
| *Firmicutes* % | 31 (21) | 58 (26) | 55 (13) | 61 (4.0) | 37 (12) | 81 (15) |
| *Proteobacteria* % | 33 (31) | 37 (28) | 2.6 (2.2) | 0.66 (0.26) | 3.6 (8.9) | 16 (13) |
| *Bacteroidetes* % | 11 (12) | 1.7 (3.9) | 28 (9.5) | 24 (3.9) | 19 (12) | 1.0 (1.9) |
| *Actinobacteria* % | 21 (15) | 0.015 (0.023) | 0.37 (0.31) | 0.69 (0.27) | 25 (16) | 1.6 (5.5) |
| *Fusobacteria* % | 0.60 (1.2) | 3.5 (5.7) | 9.6 (8.8) | 0.00 (0.00) | 5.6 (5.7) | 0.16 (0.16) |
| Shannon diversity index | 3.9 (0.84) | 2.1 (0.95) | 3.6 (0.41) | 6.5 (0.15) | 3.4 (1.1) | 1.2 (0.58) |

The microbial diversity in newborn rectal samples was relatively high (Table 2 and Fig. 4). The Shannon diversity index was significantly different between foal rectal samples collected at different time points (χ^2^= 19.7, df = 2, P = 5.2 × 10^−5^), and significantly higher in newborns compared to 24 h foals (P = 0.00024).

**FIGURE 4.**
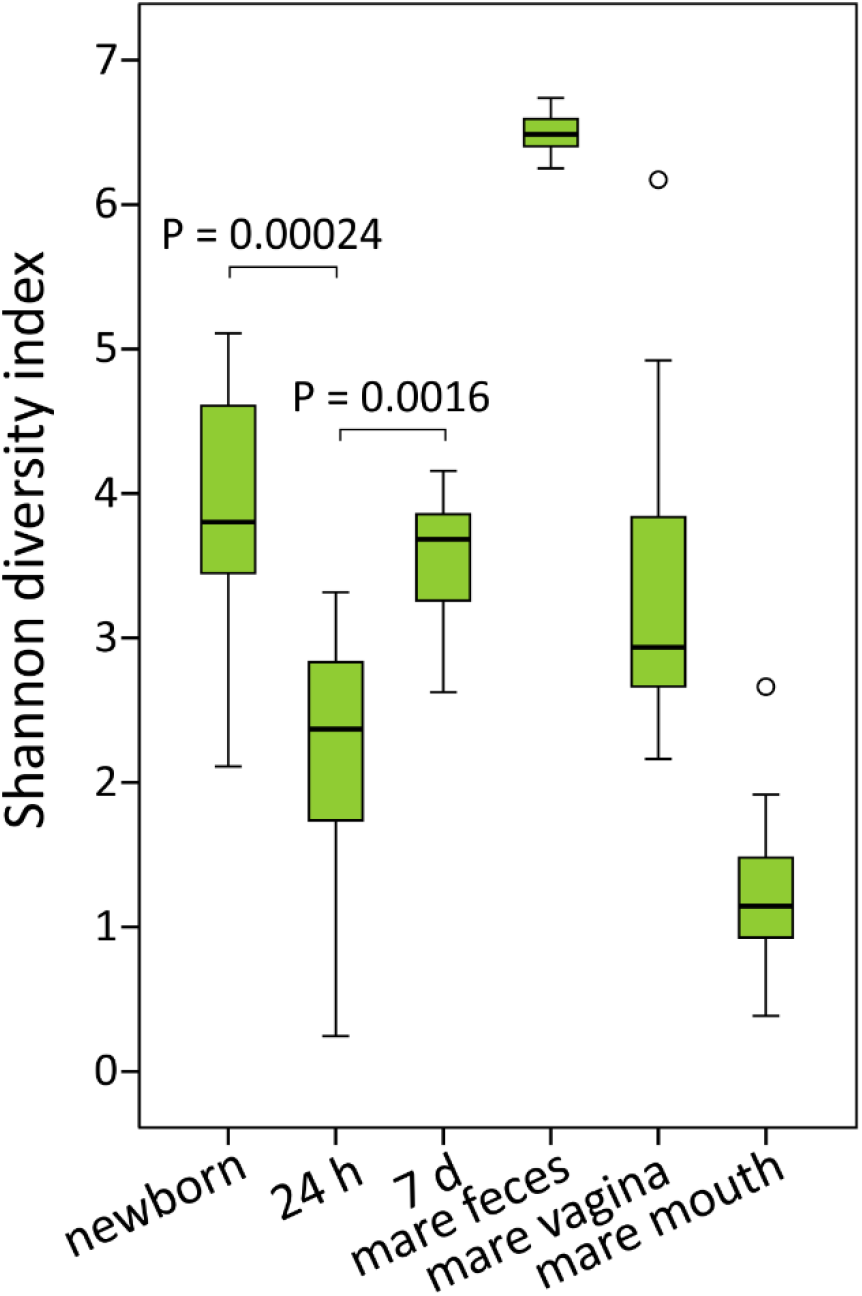
Shannon diversity indices of foal rectal samples and mare fecal, vaginal and oral samples. Boxplot as in Fig. 1.

### Postnatal development of the foal rectal microbiota

The rectal microbiota samples collected 24 hours after birth were dominated by the genus *Escherichia/Shigella* and various typical intestinal Firmicutes, especially *Clostridium* sp. (Fig. 3 and Supplementary Table 2). Members of the *Bacteroides* genus were also already detected in most of the foals at this time point. In two animals, the rectal microbiota consisted almost completely of a single genus: in one of *Escherichia/Shigella* and in the other of *Streptococcus*, with a 100% sequence match to the species *S. equinus*. The microbial diversity in the 24 h rectum was significantly lower than in the newborns (Table 2 and Fig. 4).

One week after birth, the phylum-level composition of the microbiota was already typical to adult feces (Fig. 3 and Table 2). Firmicutes was the most abundant phylum, while Proteobacteria were replaced with Bacteroidetes and Fusobacteria. *Bacteroides* was the most abundant genus. *Fusobacterium*, *Tyzzerella*, *Streptococcus* and *Lactobacillus* were also observed in all foals, and their mean relative abundance was > 5%. *Akkermansia* was detected in a majority of animals. The microbial diversity had increased in comparison to the 24 h samples (P = 0.0016) but was still clearly below the diversity of adult feces (Fig. 4).

### Mare fecal, vaginal and oral microbiota

The highly diverse mare fecal microbiota (Table 2 and Fig. 4) consisted mostly of Firmicutes and Bacteroidetes, accompanied by single genera of Kiritimatiellaeota, Spirochaetes (*Treponema*) and Fibrobacteres with mean relative abundances of 2-4% (Fig. 3 and Supplementary Table 4). At the genus level, unclassified *Lachnospiraceae*, *Rikenellaceae* RC9 gut group and several genera of *Ruminococcaceae* were most abundant (4-8% mean relative abundance).

In mare vagina, five major phyla were present in relatively equal ratios (Fig. 3). At genus level (Supplementary Table 5), the vaginal microbiota was much less diverse than the fecal microbiota (Table 2 and Fig. 4), but there were major differences between individual animals. The Firmicutes were primarily represented by six genera: *Helcococcus*, *Streptococcus, Peptoniphilus, Globicatella, Peptostreptococcus*, and *Murdochiella* (mean relative abundances 1-9%). In two mares, *Aneorococcus* was highly abundant (12% and 21%). Bacteroidetes consisted mostly of the *Porphyromonas* genus (with relative abundance varying between 0.01% and 37%), Actinobacteria of *Corynebacterium* (0.6-49%) and Epsilonbacteraeota of the more equally abundant *Campylobacter*.

The mare oral microbiota was strongly dominated by a single genus, *Gemella* from the Firmicutes phylum, which was very abundant in all animals (mean relative abundance 74%; Fig. 3 and Supplementary Table 6). The genera *Streptococcus*, *Moraxella, Enterococcus* and unclassified *Pasteurellaceae* were also detected in all mares, but their relative abundances varied between animals.

### Comparison of foal and mare microbiota compositions

We examined the differences between newborn and mare microbiotas by using principal coordinates analysis (PCoA) of unweighted Unifrac distances (Fig. 5a). All four sample types clustered separately (P = 0.001), indicating that their microbial profiles were significantly different. The newborns clustered closest to the mare vaginal microbiota. In order to compare the overall microbiota compositions in the newborns and specifically to the fecal, vaginal and oral microbiotas of their own dams, we also calculated Spearman rank correlations between these. Correlation coefficients (ρ) for all three comparisons between the newborn and mare microbiotas were between −0.002 and 0.009 (not significant).

**FIGURE 5.**
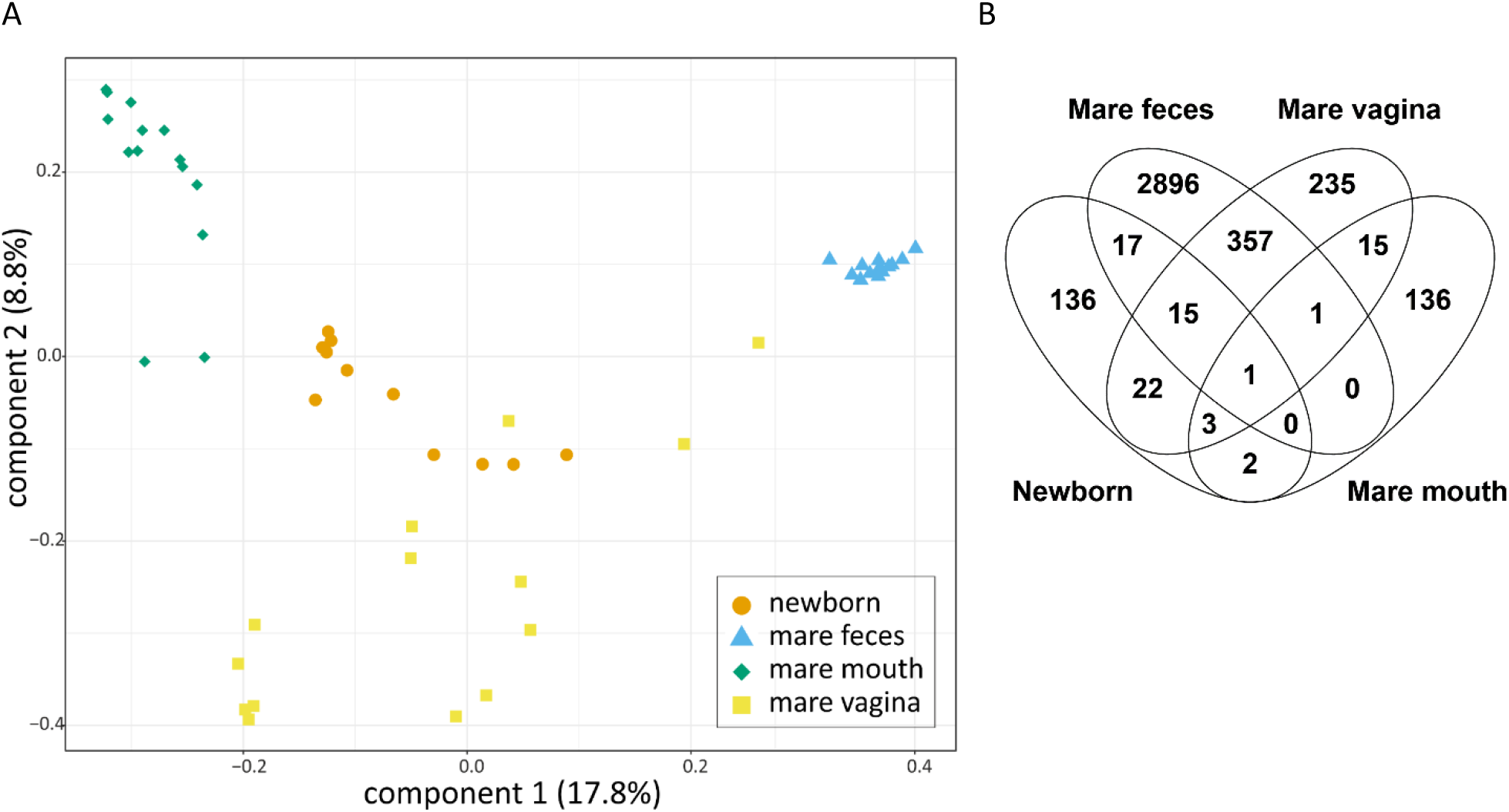
Similarity of newborn foal and mare microbiotas. A. PCoA on unweighted UniFrac distances of newborn rectal microbiota and mare fecal, oral and vaginal microbiotas from the ASV data. Colours and shapes indicate the sample types. B. ASVs shared between newborn rectal microbiota samples and various mare microbiota samples. ASVs detected in at least two animals per sample group are included.

To investigate potential similarities in the microbiotas of newborns and their own dams, we also calculated the percentages of newborn ASVs that were detected in the dam microbiotas. An individual newborn foal shared an average of 8.7% (SD = 6.9) of its ASVs with the feces of its own dam, 3.9% (SD = 3.7) with dam vagina and 0.23% (SD = 0.33) with dam mouth. These proportions of shared ASVs were significantly different (N = 8, χ^2^ = 9.9, df = 2, P = 0.0073). Newborns shared significantly more ASVs with dam feces than with dam mouth (P = 0.011). The sharing with dam vagina was not significantly different from sharing with the other microbiotas.

A Venn diagram of ASVs detected in newborns and mares as groups is shown in Fig. 5b. The most common ASVs shared between newborns and mare feces were classified as the genera *Streptococcus* (sequence match to *S. equinus*), *Ruminococcus*, *Ruminococcaceae* NK4A214 group, *Clostridium sensu stricto* 1, two different genera of *Lachnospiraceae*, *Bacteroidales* RF16 group and a genus of *Paludibacteraceae*. ASVs shared between newborns and mare vagina included *Bacillus*, *Sphingomonas*, *Campylobacter* and *Oceanivirga*. Newborns shared the most abundant *Bacillus* ASV only with mare vagina and not with mare fecal or oral microbiotas. *Sphingomonas* was shared between newborns and mare mouth. Of all the ASVs detected in any of the newborns, 38% were found in some of the mare microbiotas. On average, these comprised 42% (SD = 19) of sequence reads in the individual newborns.

At 24 hours, the foal rectal microbiota clustered separately from all mare microbiotas in PCoA (not shown), and no significant correlations between the 24 h foal and mare microbiotas were observed. On average, the 24 h foal samples shared 5.2% (SD = 4.7) of their ASVs with the vagina of their own dam, 1.7% (SD = 2.3) with dam feces and 1.5% (SD = 2.5) with dam mouth. These differences were significant (N = 12, χ^2^ = 8.4, df = 2, P = 0.015). The 24 h foals shared significantly more ASVs with the vagina of their own dam than with the dam mouth (P = 0.05). The proportion of ASVs shared with dam feces did not differ significantly from the proportions shared with the other dam microbiotas.

A Venn diagram of ASVs observed in the 24 h foals and mare microbiotas is shown in Supplementary Fig. 3a. The foals shared abundant ASVs classified as *Clostridium sensu stricto 1*, *Bacteroides*, *Terrisporobacter* and *Fusobacterium* exclusively with vagina. Of all the ASVs detected in any 24 h foal, 29% were observed in some of the mare microbiotas. On average, these comprised 47% (SD = 24) of sequence reads in the individual foals.

The 7-day rectal microbiota still clustered separately from all mare microbiotas (not shown), with no significant correlations between different mare microbiotas. However, the phylum-level composition resembled the adult fecal microbiota (Fig. 3 and Table 2), although some of the adult phyla were still not detected in all or most foals (Kiritimatiellaeota, Spirochaetes, Fibrobacteres and several low-abundance phyla such as Patescibacteria, Lentisphaerae and Cyanobacteria). At family level, the day 7 microbiota clustered close to the adult fecal microbiota in PCoA (Supplementary Fig. 4).

The 7 d foals shared an average of 3.3% (SD = 2.8) of their ASVs with the feces of their own dams, 2.8% (SD = 2.0) with dam vagina and 1.3% (SD = 1.7) with dam mouth. The differences in proportions of shared ASVs were significant (N =14, χ^2^ = 6.5, df = 2, P = 0.039). The 7 d foals shared significantly more ASVs with dam feces than with dam mouth (P = 0.037). The sharing with dam vagina did not differ significantly from the sharing with the other dam microbiotas. A Venn diagram of ASVs detected in 7 d foals and mare microbiotas is shown in Supplementary Fig. 3b. *Bacteroides* and *Fusobacterium* ASVs were exclusively shared with mare vagina. Out of all detected ASVs in the 7 d foals, 28% were observed in some of the dam microbiotas. These comprised an average of 38% (SD = 13) of reads in the individual foals.

### The change of rectal microbiota composition over time

The fecal microbiota of all four age groups (newborns, 24 h foals, 7 d foals and adult mares) clustered separately from each other in PCoA (Fig. 6a), indicating that the microbiota compositions were significantly different (P = 0.001).

**FIGURE 6.**
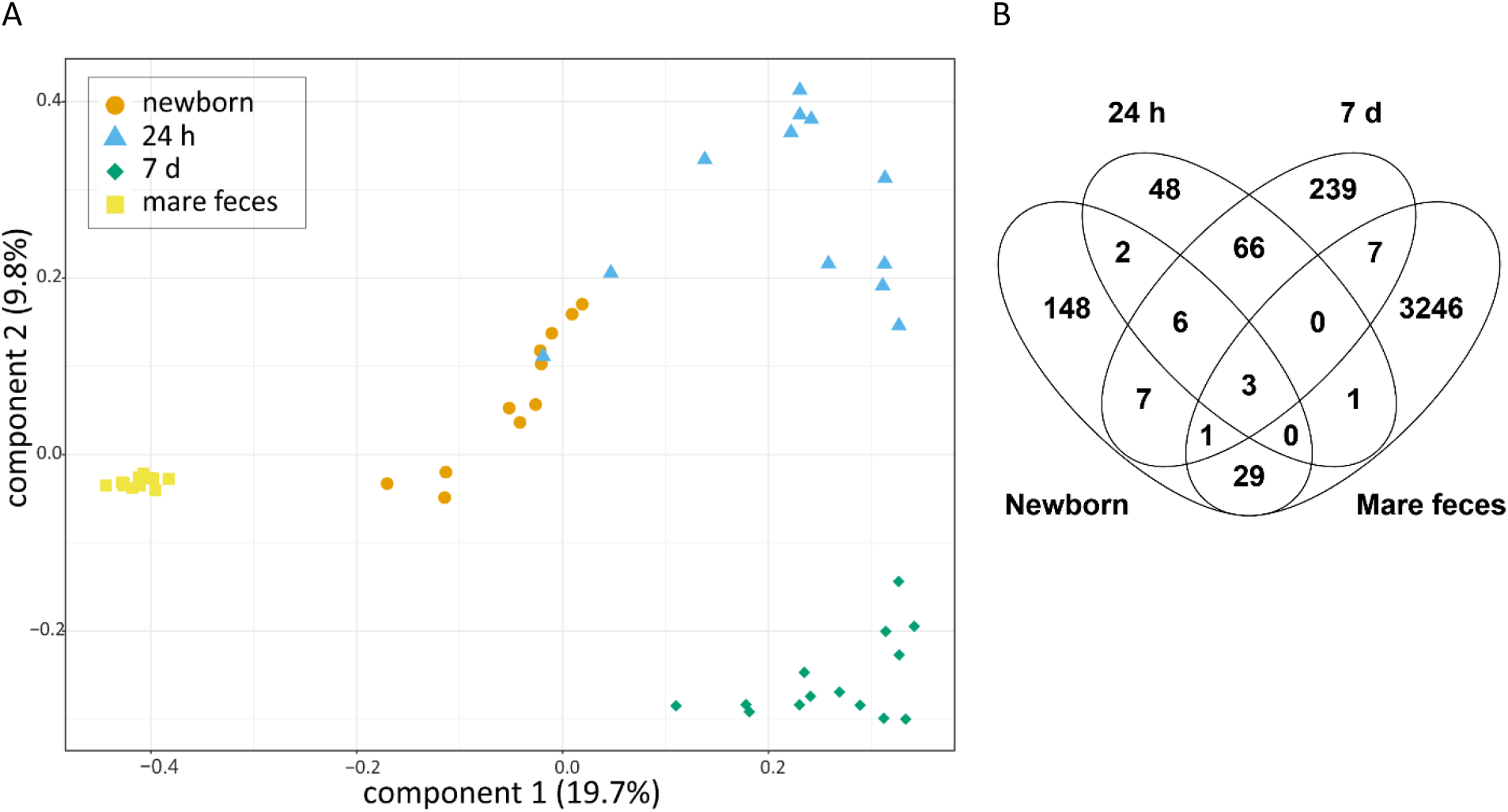
Similarity of horse fecal microbiotas at various ages. A. PCoA on unweighted UniFrac of rectal microbiota in the newborn, 24 h old and 7 d old foals and adult mares from ASV data. Colours and shapes indicate the sample types. B. ASVs shared between rectal microbiota samples at various ages. All ASVs detected in any animal are included. Therefore, the numbers are larger than the mean ASV counts presented in Table 1.

The newborn rectal samples grouped closest to the 24 h fecal samples in PCoA. However, no significant correlation was observed between newborns and the same foals at 24 h (N = 6, average ρ = 0.03, SD = 0.04, P = 1), confirming that the overall microbiota compositions were different. An average of 2.9% (SD = 4.1) of newborn ASVs was detected at 24 hours in the same foal. A Venn diagram of ASVs observed in each age group is shown in Fig. 6b. The ASVs shared between newborns and 24 h foals as groups were classified to the genera *Escherichia/Shigella*, *Epulopiscium*, *Fusobacterium*, *Streptococcus* and *Butyricicoccus*.

A significant correlation was observed between the 24 h foals and the same foals at day 7 (N = 12, average ρ = 0.21, SD = 0.11, P = 0.001). Of the ASVs observed at 24 hours, an average of 31% (SD = 12) was detected in the same foals at day 7. The most common of these included ASVs classified to the genera *Tyzzerella 4*, *Faecalitalea*, *Clostridium sensu stricto 1*, *Fournierella* and *Erysipelatoclostridium*.

The phyla Actinobacteria and Verrucomicrobiota were not yet detected in most foals at 24 h, although they were present in all or most of them at day 7. Of the relatively common ASVs, only one classified as *Streptococcus equinus* was found in the rectal microbiota at all ages, from newborns to adults.

## Discussion

In this study, we observed the early development of the equine intestinal microbiota over the first week of postnatal life, starting from the moment of birth. In order to determine the potential origins of the foal microbiota, we characterized the vaginal and oral mucosal microbiota of the mares, which have not been previously investigated by sequencing, as well as the mare fecal microbiota.

In order to maximize the reliability of the microbiota analyses, we applied several types of negative controls and performed a stringent filtering of the sequencing data to remove potential contaminants. Our data decontamination protocol was based on a previously published logic, comparing the prevalence and relative abundance of each taxon in samples and negative controls^27,31,32^. We performed filtering on the amplicon sequence variants (ASVs), which provides maximum resolution of the 16S rRNA gene sequencing data^29,30^. This is important, as filtering at a too coarse taxonomic level causes false negatives by deleting related but different taxa observed in samples and controls. We set stringent criteria for ASV acceptance to minimize false positive observations. Experimenting with varying filtering thresholds had little effect to the proportion of discarded ASVs, indicating that this did not cause a large number of false negatives. ASVs classified as typical intestinal genera, which are unlikely reagent contaminants were well preserved, while known reagent contaminants such as *Ralstonia* were efficiently removed. The abundance of likely contaminants in the unfiltered raw data underlines the importance of careful controls and data decontamination in the analysis of sensitive microbiota samples.

In majority of the newborn foals, the rectum contained a small amount of bacterial DNA, which was detectable with quantitative PCR and distinguishable from negative controls by sequencing. Bacterial DNA was also detected in a recently published study of foal meconium, but the protocol did not include negative controls or quantification, which are necessary for reliable analyses of low-abundance microbiotas^24^. Almost all of the Firmicutes genera observed in our newborns, as well as the most prevalent genera of other phyla, represented typical mucosal taxa and were shared with the core vaginal and/or fecal microbiota of the mares. Several of these genera were reported in the previous analysis of newborn foal microbiota and many were also observed by us in newborn calf rectum^24,27^. The prevalent ASVs with species-level classification by our analysis pipeline showed sequence matches to equine-associated species. These observations indicate that the rectum in these foals indeed harboured bacterial DNA already at the moment of birth. These probably represent the late fetal status, as rectal colonization during the equine parturition is unlikely. The foals were sampled immediately after birth and the outer chorioallantoic membrane typically ruptures only 10-30 minutes before the foal is born. The inner amniotic membrane covers the newborn completely or at least from neck to rump.

Previous studies in bovine calves^27^, mice^28^ and humans^33,34^ have suggested that the fetus may be seeded with microbes from the mouth or proximal digestive tract of the mother. In our newborn foals, the rectal microbiota was more similar to the dam vaginal and fecal microbiotas than to the dam oral microbiota. This may reflect species differences in the digestive physiology. In ruminants, the forestomach harbours the microbiota essential for forage fermentation; in horses, it resides in the large intestine. Of all the ASVs detected in any newborn, 62% were not detected in any of the three mare microbiotas analysed, suggesting that they originated from other sources. Within the digestive tract, small intestine, colon and caecum are more likely sources for transplacental microbial translocation than rectum, as the luminal microbiota is sampled by the immune system in these locations^35^.

At 24 hours, the rectal microbiota contained a large proportion of *Escherichia/Shigella*, as previously observed in calves^27^. In contrast to 24 hour old calves, most foals were already also colonized by *Bacteroides* and several genera of the Firmicutes, which are common in intestinal microbiota. The foals shared most ASVs with dam vagina, and several abundant ASVs were shared exclusively with vagina. This suggests that microbial colonization during birth is significant, although the resolution of the 16S rRNA gene sequencing is not sufficient to prove direct microbial transfer. Interestingly, one of the most prevalent Firmicutes in the 24 hour samples was classified as the genus *Epulopiscium*, previously known from a giant bacteria of the surgeonfish gut^36^. These ASVs probably represent uncharacterized members of mammalian *Lachnospiraceaea*, as identical 16S rRNA sequences are present in a human ileal metagenome^37^.

Previous studies have reported different compositions for 24 h old foal rectal microbiota. Costa et al. reported a larger proportion of Firmicutes and less Proteobacteria; their foals had much less *E. coli* and more Ruminococcaceae, Lachnospiraceaea and *Lactobacillus*^25^. In the study by De La Torre et al., first-day foal microbiota was mostly composed of *Acinetobacter* and other *Moraxellaceae*, which were rare in our 24 h foals^23^. This difference may be due to the exact timing of sampling of the rapidly changing early microbiota: De La Torre et al. sampled the foals at variable times during the first postnatal day. The early microbiota is probably also affected by the environment. In our study, the foals were allowed in outdoor paddocks from the beginning, whereas in the study by De La Torre and colleagues the foals were first kept mostly in stalls.

Several genera commonly observed as cause for neonatal foal sepsis were prevalent in the 24 h rectal samples, including *Escherichia/Shigella, Streptococcus, Enterococcus* and *Klebsiella*^38,39^. The rectal microbiota of one foal consisted almost completely of *Escherichia/Shigella*, and that of another foal of ASVs classified as *Streptococcus equinus*. No obvious adverse effects were observed in these foals, and at day 7 their rectal microbiota was similar to the other foals. A dominance of these bacteria in the gut at 24 hours of age is not necessarily problematic to the host.

At day 7, more than 30% of ASVs observed in the same foal at 24 hours were still detectable, but the overall composition was markedly different. At the phylum level, the day 7 rectal microbiota was already very similar to the adult mares, and clustered close to the adults also at the family level. However, the core genera were largely distinct, and only 3% of ASVs in the day 7 rectum were detectable in the feces of the foal’s own dam. Only an ASV classified as *Streptococcus equinus* was common in the rectum at all ages. Foals typically eat their dam’s manure^40^, but colonization is probably restricted by differences in diet and intestinal environment. Similar observations were recently reported by De La Torre *et al.*: the foal microbiota was very different from the adults at day 7, and developing towards the adult composition at day 28^23^. Their day 7 gut microbiota was similar with our observations at the family level. The most abundant genus in our 7 d foals, *Bacteroides*, was found almost exclusively in the vaginal microbiota in the mares. Thus, the microbial contribution of the birth canal appears to be quite stable in the young foals. Vagina was recently reported as a source of physiologically important rumen microbes in calves^9^.

We could trace 28-29% of all ASVs observed in 24 h and 7 d foals to the three mare microbiotas analysed. These shared ASVs comprised 38-48% of sequence reads in the foals. This suggests that the foals obtained a majority of their microbiota from other sources. Some of the foal microbes could be present at undetectably low relative abundances in the mare samples, even though we performed relatively deep sequencing. Milk and dam skin microbiotas are obvious contributors which we did not analyse. In the previous study by Quercia *et al*.^24^, the foal intestinal microbiota at 0-3 days after birth was found to resemble the dam milk microbiota.

In our mares, the composition of the fecal microbiota was largely similar to that recently reported for pregnant horses^23^. The relative abundances of the *Lachnospiraceae* and *Rumonicoccaceae* families and the order Bacteroidales were similar, but in our mares, *Prevotellaceae* were much more abundant and various Clostridiales less abundant. In contrast, the fecal microbiota composition in non-pregnant horses was different even at the phylum level: fecal Firmicutes were less abundant and Bacteroidetes more abundant^41,42^. This may be due to the supplemented feed of pregnant mares, or other effects of pregnancy. Kirimatiellaeota were not previously reported in equine feces, probably because this novel phylum was not included in the taxonomical databases used in those studies. The most abundant streptococci (classified as sp. 27284-01) in mare feces were different from the dominant species in the vaginal and foal rectal microbiotas.

To our knowledge, the healthy vaginal microbiota of pregnant mares has not been previously characterized. Compared to cows, the vaginal microbiota in our mares was more distinct from the fecal microbiota, as the mare vagina is typically less contaminated with feces^27^. *Corynebacterium*, *Porphyromonas*, *Helcococcus* and *Campylobacter* were the most abundant genera. Most of these have been previously observed in the bovine vaginal microbiota, where the abundance of *Helcococcus* is associated with metritis^43,44^.

Interindividual differences were greater for the vaginal than the fecal microbiota. In contrast to humans, very few lactobacilli were detected in the mare vagina, as observed previously in a culture based study^45^. This is probably associated with the higher pH of the mare vagina.

The microbiota of the oral vestibular mucosa in horses has also not been previously investigated. In our mares, the phylum-level composition was more similar to human buccal microbiota than to bovine oral microbiota, which is affected by rumination^46,47^. However, Bacteroidetes were less abundant in the horses than in the human. At the genus level, the mare oral microbiota was strongly dominated by *Gemella*, whereas in human mouth streptococci are typically most abundant. *Pasteurellaceae*, *Neisseria* and *Porphyromonas* are found in both species. Gemella and the less frequently observed *Actinobacillus* have previously been associated with good dental health in equine subgingival samples^48^.

In conclusion, bacterial DNA was detected in the foal intestine at birth, suggesting it was present already prenatally. The microbial profile at birth was closer to mare vaginal and fecal microbiota than mare oral microbiota. The rectal microbiota changed rapidly. At one week of age, the phylum-level composition already resembled adult fecal microbiota, but the bacterial genera were different. The impact of the mare vaginal microbiota was still detectable at day 7. The pregnant mare fecal microbiota was different from previously studied nonpregnant horses, due to feeding or other effects of pregnancy. The mare vagina contained few lactobacilli, and the oral mucosal microbiota was dominated by *Gemella* rather than streptococci.

## Methods

### Animals and sample collection

Microbiota samples were collected from fourteen standardbred mare-foal pairs and four additional foals in the spring foaling seasons 2016-2017. Two mares were included in both seasons of the study. Mares were 6 to 16 years old with a parity of 1 to 6. Foals came from 15 different sires. There were 11 fillies and 8 colts.

Ultrasonographic monitoring of fetal-placental wellbeing was performed one month before foaling and all mares were diagnosed as clinically normal. The mares received an influenza and tetanus vaccination one month before parturition. They were not treated with antibiotics or other medications during the last three months of the pregnancy. All mares lived in same barn with 32 stalls. The barn was cleaned, washed and disinfected between breeding seasons. Mares were housed in separate wide straw-bedded stalls, fed analysed high quality haylage ad libitum and 2 liters of Krafft Groov protein concentrate (Lantmännen Krafft Ab, Sweden; digestible raw protein 115 g/kg) twice per day.

Parturitions took place between March and June. Every mare had a Birth Alarm foaling belt (Birth Alarm, UK) and a camera in their stall to alert at the start of parturition. Before parturition, mares were kept outside during the day in groups of four. One or two days after parturition, mare-foal pairs were kept short time periods every day in small individual paddocks. The mean length of the pregnancy was 333 ± 8 d (normal gestation period 340 d) and all the parturitions were uneventful.

All foals were assessed clinically healthy by a veterinarian and received colostrum within first 1.5 hours. The concentration of blood IgG was measured using a quantitative immunoassay when the foals were 24 - 36 hours old. All foals had adequate IgG concentrations.

Sample collection was performed by the same veterinarian for all the animals. Foal samples were collected from within the rectum (5-10 cm from anus) using sterile double sheathed Minitube uterine culture swabs (Minitüb, Germany) while wearing sterile gloves, as described previously^27^. Newborn samples were collected within 10 minutes after birth. Mares were not allowed to lick the foals before sample collection. The foal sampling was repeated 24 h after birth and 7 d after birth. Field controls were collected by exposing Minitube swabs to air in the sampling environment. Mare samples were collected before foaling with sterile cotton sticks from feces (from the inside of fecal balls retrieved from rectum), labial part of the lower vestibulum oris (behind the lower lip) and ventral side of the vaginal vestibulum before foaling. Samples were kept at −25 °C for a short period of time, before storing at −80 °C.

The experimental protocols were reviewed and approved by the University of Helsinki Viikki Campus Research Ethics Committee. The sampling was carried out in accordance with animal welfare guidelines and regulations.

### DNA extraction

DNA from the sampling swabs was extracted using the ZymoBIOMICS™ DNA Miniprep Kit (Zymo Research, Irvine, CA, USA) according to the manufacturer’s instructions with minor modifications to the bead beating process, as described here. The bead beating was done using a FastPrep^®^ 24 Instrument (MP Biomedicals, Inc., USA) in two successive rounds at 5.5 m/s for 3 minutes. In order to prevent excessive shearing of DNA, the lysate fraction following the first round bead beating was collected and additional 200 µl ZymoBIOMICS™ Lysis Solution was added before performing the second round. The lysate produced in the second bead beating round was pooled with the previous lysate prior to continuing the procedure according to the manufacturer’s instructions.

Unopened (instrument controls, N = 6) and opened (field controls, N = 6) empty uterine culture swabs were used as negative controls and processed in the same batch with the newborn samples. The other, higher-biomass samples were processed separately to avoid cross-contamination to the newborn samples. ZymoBIOMICS™ Microbial Community Standard was processed with the high-biomass samples. No-template-controls were included in every batch. The phases involving opening the tubes were performed in laminar flow cabinet, and the workplace, instruments and pipettes were cleaned routinely with 10 % bleach. Certified DNA, RNase, DNase and PCR inhibitor free tubes (STARLAB International, Germany) and Nuclease-free Water (Ambion™, Thermo Fisher Scientific, USA) were used for DNA extraction and downstream analyses. Extracted DNA was stored at −80 °C.

### Quantitative PCR

The absolute 16S rRNA gene copy numbers in the DNA extracted from the foal rectal samples and the controls was determined using quantitative PCR as described previously^27,49^ and in Supplementary Methods. Universal eubacteria probe and primers were used in the amplification.

### Library preparation and 16S rRNA gene amplicon sequencing

The hypervariable region V3-V4 of 16S rRNA gene was sequenced using the Illumina MiSeq platform in the DNA core facility of University of Helsinki, as described previously^50^ and in Supplementary Methods.

To compensate for different quantities of 16S rRNA gene copies in the samples the newborn meconium samples, instrument controls and field controls were preamplified with 21 PCR cycles, mare mouth and vaginal samples with 18 cycles, 24 h foal rectal samples with 16 cycles and 7 d old foal rectal samples and mare fecal samples with 14 cycles. ZymoBIOMICS™ Microbial Community Standard (Zymo Research, U.S.A.) was preamplified with 12 cycles.

### Bioinformatics

The detailed bioinformatics pipeline is reported in the Supplementary Methods. Briefly, remaining primer and spacer sequences were first removed with Cutadapt v.1.10^51^. The paired reads were imported into QIIME2 v2018.8^52^. The sequences were truncated to remove low quality regions. The DADA2 pipeline plugin in QIIME2 was used to model and recognize the amplicon sequencing errors in the Illumina sequencing data and to infer the exact sample sequences^30^. This resulted in an amplicon sequence variant (ASV) table. For the taxonomic analysis, curated taxonomy was extracted from the SILVA v132 QIIME release 97% database and a naïve Bayes classifier was trained based on it^53^. The classification was completed with the q2-feature-classifier plugin using scikit-learn^54^.

ASVs that could not be assigned a taxonomic classification based on the SILVA database, unclassified bacteria (mostly mitochondrial sequences), chloroplasts and the ASVs which were detected less than 10 times in the entire dataset were removed. The data was then filtered to remove ASVs which represented probable contaminants. An ASV was removed if its prevalence in actual samples was ≤ 2× its prevalence in instrument controls *or* its prevalence in field controls, *or* if its mean relative abundance in actual samples was ≤ 10× its mean abundance in instrument controls *or* its mean abundance in field controls. The filtering was performed for each sample group separately.

### Statistics

Mann-Whitney U test was used to assess the statistical significance of the difference between qPCR measurements in newborns and field controls using IBM SPSS Statistics 23.Shannon diversity indices were calculated for ASV data with the R package phyloseq^55^. Kruskal-Wallis rank sum test was used to compare Shannon diversities between newborn, 24 h and 7 d foal samples with base R^56^. Pairwise post-hoc comparisons were then performed with Nemenyi’s all-pairs test with χ^2^ approximation, using the R package PMCMRplus^57^.The PCoA figures were plotted using ASV data and unweighted UniFrac distances with the R package phyloseq^55^. Permutational analysis of variance (PERMANOVA) and multivariate homogeneity of groups dispersions (betadisper) were calculated using the R package vegan^58^.The percentages of shared ASVs were compared between sample groups using Friedman rank sum test with unreplicated blocked data using R. Pairwise comparisons were performed using Nemenyi’s all-pairs comparison test for unreplicated blocked data using the R package PMCMRplus. The Venn diagrams were generated using Venny^59^. Spearman rank correlations were calculated to compare newborn foals and their own dams, as well as each foal at different time points, using the R package vegan^58^. The P values were Bonferroni corrected for multiple testing.

### Data availability

The raw 16S rRNA gene amplicon sequencing dataset is available in the European Nucleotide Archive with the ENA accession number PRJEB32017.

## Acknowledgements

We thank Kirsi Lahti, Santeri Suokas and Oskari Mäenpää for expert technical assistance, and Anna Mykkänen for commenting the manuscript. The study was funded by University of Helsinki and the Finnish Veterinary Foundation. JJ was supported by a fellowship from the Academy of Finland (grant no. 0313471-7).

## Author contributions

AH performed the PCR amplification for sequencing, organized the amplicon sequencing, processed and analysed the data and drafted the manuscript; JJ participated in the data analysis and interpretation of results; MJA performed most of the DNA extractions; PH participated in study design and collected the samples; MK participated in study design; AI and TPM participated in study design and data analysis; MN conceived, designed and organized the study and participated in data processing and analysis. All authors participated in manuscript preparation and proved the final draft.

## Additional information

The authors declare the following competing interests: PH is an owner of Sahara stud.

## Notes

https://www.ebi.ac.uk/ena/data/view/PRJEB32017

