## Supplementary_info for "The composition of the perinatal intestinal microbiota in horse"

### Supplementary information for Husso et al. 2019:
### The composition of the perinatal intestinal microbiota in horse

Husso A[1], Jalanka J[2], Alipour MJ[1], Huhti P[3], Kareskoski M[4], Pessa-Morikawa T[1], Iivanainen A[1*], Niku M[1*]

[1] Veterinary Biosciences, Faculty of Veterinary Medicine, University of Helsinki, Finland

[2] Human Microbiome Research Program, Faculty of Medicine, University of Helsinki, Helsinki, Finland

[3] Saharan ravitalli (Sahara stud), Haapamäki, Finland

[4] Production Animal Medicine, Faculty of Veterinary Medicine, University of Helsinki, Finland

Antti Iivanainen and Mikael Niku jointly supervised this work.

### Supplementary results and discussion

#### *Observed composition of the commercial microbiota community standard*

The ZymoBIOMICS Microbial Community Standard (Zymo Research, Irvine, CA, USA) was used to assay the reliability of the entire analytical pipeline from DNA extraction in representing the actual microbiota community composition. A comparison of the observed and expected compositions and taxonomic classifications are shown in Fig. S1. The expected 16S rRNA gene composition is based on the data provided by the manufacturer.

The observed composition closely matched the expected composition. The relative abundances of *Pseudomonas aeruginosa, E. coli* and *Bacillus subtilis* were slightly overestimated, while those of *Listeria monocytogenes* and *Lactobacillus fermentum* were slightly underestimated. *Listeria monocytogenes, Lactobacillus fermentum* and *Staphylococcus aureus* were correctly classified to the species level. A minority of *S. aureus* sequences and all the other bacteria were correctly classified to the genus level.

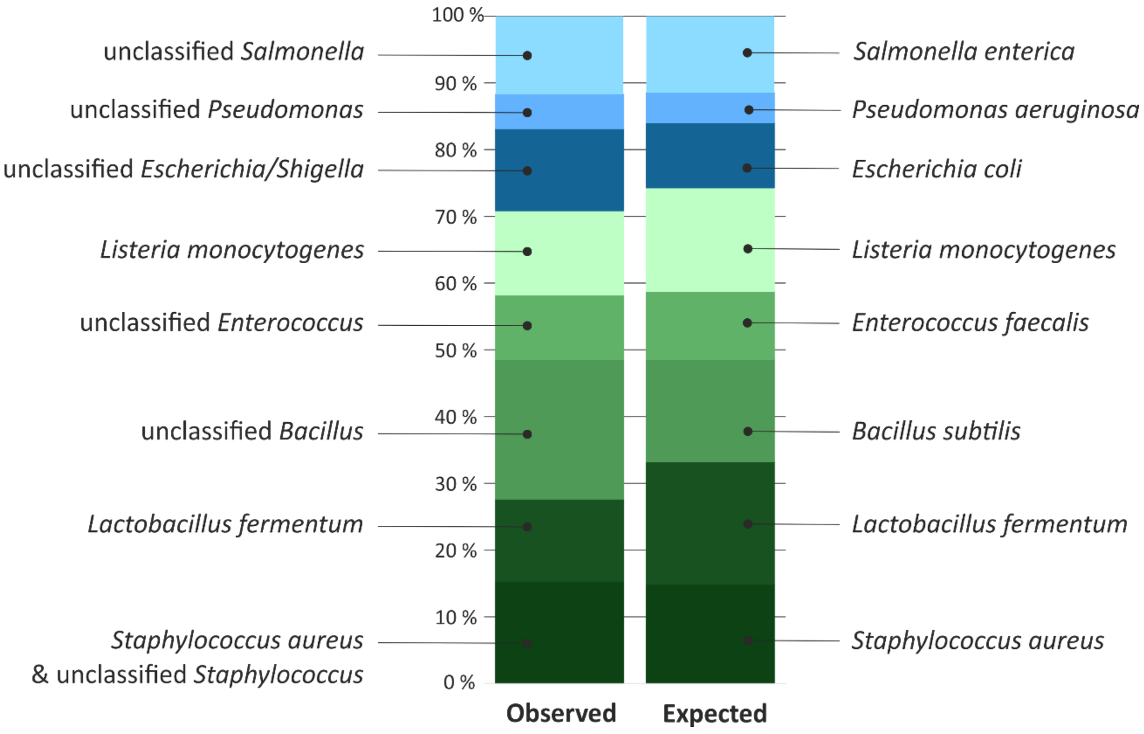

***SUPPLEMENTARY FIGURE 1.*** *Observed (left) and expected (right) relative abundances of the bacteria included in the commercial community standard. Blue = Proteobacteria, green = Firmicutes.*

##### *Effect of data decontamination on phylum-level microbiota compositions and read counts*

Average phylum-level compositions and read counts in unfiltered raw data and in decontaminated data are shown in Supplementary Fig. 2. Almost all ASVs detected in negative controls were removed. In newborns, majority of data was also removed as potential contaminants, but in 11 out of 18 animals, a 16 rRNA gene profile could be observed and characterized. In samples from older animals, the filtering had minor effects on average microbiota compositions and read counts.

The filtering effectively removed obvious reagent contaminants such as Ralstonia (see Results). To estimate the rate of false negatives, we calculated the proportion of reads deleted from several typical intestinal taxa. These included ASVs classified as *Akkermansia, Bacteroides, Christensenellaceae, Clostridium sensu stricto 1, Enterococcus, Faecalibacterium, Fibrobacter, Lachnospiraceae, Lactobacillus, Prevotella, Ruminiclostridium* and *Ruminococcaceae*, as well as ASVs labelled as “gut group” or “gut metagenome”. As described in Results, only 0.18% of these were purged.

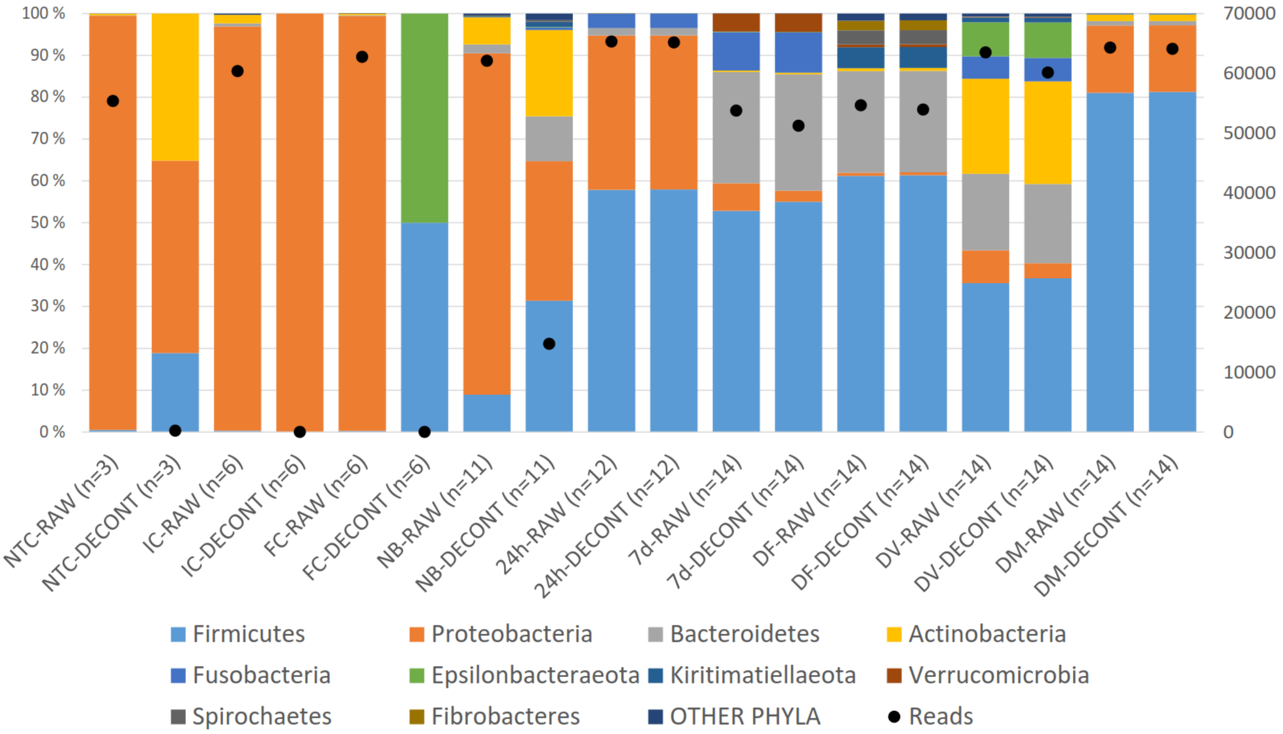

***SUPPLEMENTARY FIGURE 2.*** *Effect of data decontamination on average phylum-level compositions and read counts. NTC = PCR no-template control; IC = instrument control; FC = field control; NB = newborns; 24h = 24h old foals; 7d = 7d old foals; DF = dam feces; DV = dam vagina; DM = dam mouth; RAW = unfiltered raw data; DECONT = decontaminated data. Newborn foal samples with less than 1500 accepted ASVs and two low-quality 24h samples were excluded from the analysis.*

#### *Core bacterial taxa in foals*

Most prevalent genus-level bacterial taxa (present in >50% of animals) in newborn and older foals are shown in Supplementary Tables 1 – 3.

***SUPPLEMENTARY TABLE 1: Genus-level bacterial taxa observed in newborn foal rectum in >50% of animals.***

*Prevalences, mean relative abundances and standard deviations in newborn foals (n=11), and sharing of taxa with the other samples (FE = mare feces, VA = mare vagina, MO = mare mouth).*

| ***Taxon*** | ***Prevalence*** | ***Mean*** | ***SD*** | ***24h*** | ***7d*** | ***FE*** | ***VA*** | ***MO*** |
|---|---|---|---|---|---|---|---|---|
| **Actinobacteria** | **91 %** | **20.65 %** | **14.61 %** | | | | | |
| *Corynebacterium 1* | 82 % | 3.43 % | 3.58 % | - | - | - | + | - |
| *Brachybacterium* | 64 % | 2.77 % | 4.30 % | - | - | - | - | - |
| unclassified *Microbacteriaceae* | 64 % | 2.04 % | 4.02 % | - | - | - | - | - |
| *Nocardioides* | 64 % | 0.47 % | 0.68 % | - | - | - | - | - |
| *Kocuria* | 55 % | 0.90 % | 1.92 % | - | - | - | - | - |
| *Glutamicibacter* | 55 % | 0.43 % | 0.91 % | - | - | - | - | - |
| *Streptomyces* | 55 % | 0.40 % | 0.75 % | - | - | - | - | - |
| **Bacteroidetes** | **100 %** | **10.70 %** | **12.43 %** | | | | | |
| *Chryseobacterium* | 73 % | 0.77 % | 1.08 % | - | - | - | - | - |
| *Pedobacter* | 64 % | 1.56 % | 2.50 % | - | - | - | - | - |
| *Rikenellaceae RC9* gut group | 64 % | 0.75 % | 1.23 % | - | - | + | + | - |
| *Sphingobacterium* | 55 % | 0.58 % | 1.13 % | - | - | - | - | - |
| *Hymenobacter* | 55 % | 0.31 % | 0.72 % | - | - | - | - | - |
| *Bacteroides* | 55 % | 0.21 % | 0.37 % | + | + | - | + | - |
| **Firmicutes** | **100 %** | **31.38 %** | **20.90 %** | | | | | |
| *Staphylococcus* | 91 % | 8.77 % | 12.61 % | - | - | - | + | - |
| *Bacillus* | 82 % | 2.20 % | 2.78 % | - | - | - | + | - |
| *Streptococcus* | 82 % | 1.51 % | 1.28 % | + | + | + | + | + |
| unclassified *Lachnospiraceae* | 73 % | 0.71 % | 0.91 % | + | + | + | + | - |
| *Lactobacillus* | 64 % | 3.23 % | 9.87 % | - | + | + | + | - |
| *Clostridium sensu stricto 1* | 64 % | 0.50 % | 0.74 % | + | + | + | + | - |
| *[Eubacterium] coprostanoligenes* group | 55 % | 0.92 % | 2.15 % | - | + | + | - | - |
| *Ruminococcaceae UCG-010* | 55 % | 0.79 % | 1.60 % | - | - | + | + | - |
| *Anaerovorax* | 55 % | 0.40 % | 0.76 % | - | - | + | - | - |
| *Ruminococcaceae UCG-005* | 55 % | 0.37 % | 0.67 % | - | - | + | - | - |
| *Paenibacillus* | 55 % | 0.20 % | 0.28 % | - | - | - | - | - |
| *Phascolarctobacterium* | 55 % | 0.15 % | 0.28 % | - | + | + | - | - |
| **Kiritimatiellaeota** | **55 %** | **1.17 %** | **2.18 %** | | | | | |
| uncultured *WCHB1-41* | 55 % | 0.84 % | 1.49 % | - | - | + | - | - |
| **Proteobacteria** | **100 %** | **33.35 %** | **30.48 %** | | | | | |
| *Ralstonia* | 100 % | 1.42 % | 1.43 % | - | - | - | - | - |
| *Sphingomonas* | 91 % | 1.74 % | 1.78 % | - | - | - | + | - |
| *Pseudomonas* | 82 % | 1.13 % | 1.79 % | - | - | - | - | - |
| *Methylobacterium* | 64 % | 5.25 % | 10.66 % | - | - | - | - | - |
| *Acinetobacter* | 64 % | 1.65 % | 2.16 % | - | - | - | - | - |
| *Paracoccus* | 64 % | 0.68 % | 0.78 % | - | - | - | - | - |
| *Allorhizobium-Neorhizobium-Pararhizobium-Rhizobium* | 55 % | 2.00 % | 3.36 % | - | - | - | - | - |
| *Bradyrhizobium* | 55 % | 1.62 % | 2.21 % | - | - | - | - | - |

***SUPPLEMENTARY TABLE 2: Genus-level bacterial taxa observed in 24 h old foal rectum in >50% of animals.***

*Prevalences, mean relative abundances and standard deviations in 24 h old foals (n=12), and sharing of taxa with the other samples (NB = newborn, FE = mare feces, VA = mare vagina, MO = mare mouth).*

.

| *Taxon* | *Prevalence* | *Mean* | *SD* | *NB* | *7d* | *FE* | *VA* | *MO* |
|---|---|---|---|---|---|---|---|---|
| **Bacteroidetes** | **75 %** | **1.73 %** | **3.86 %** | | | | | |
| *Bacteroides* | 75 % | 1.67 % | 3.73 % | - | - | - | - | - |
| **Firmicutes** | **100 %** | **57.96 %** | **26.20 %** | | | | | |
| *Streptococcus* | 92 % | 9.42 % | 28.57 % | + | + | - | + | - |
| *Epulopiscium* | 92 % | 4.24 % | 4.69 % | | | | | |
| *Clostridium sensu stricto 1* | 83 % | 24.04 % | 18.68 % | | | | | |
| *Terrisporobacter* | 83 % | 5.66 % | 4.52 % | + | + | + | + | + |
| *Romboutsia* | 83 % | 3.73 % | 5.03 % | - | - | - | - | - |
| *Enterococcus* | 83 % | 0.66 % | 0.72 % | + | + | + | + | - |
| *Tyzzerella 4* | 75 % | 1.57 % | 4.30 % | - | + | - | - | - |
| *Faecalitalea* | 75 % | 0.90 % | 1.42 % | - | - | - | - | - |
| *[Ruminococcus] gnavus* group | 75 % | 0.50 % | 0.52 % | - | - | - | - | + |
| *Erysipelatoclostridium* | 75 % | 0.15 % | 0.17 % | - | + | - | - | - |
| unclassified *Lachnospiraceae* | 67 % | 1.94 % | 2.97 % | - | + | - | - | - |
| *Turicibacter* | 67 % | 1.40 % | 2.48 % | - | + | - | - | - |
| *Butyricicoccus* | 67 % | 0.67 % | 0.93 % | - | + | + | - | - |
| *Fournierella* | 58 % | 0.94 % | 1.56 % | + | + | + | + | - |
| unclassified *Ruminococcaceae* | 58 % | 0.12 % | 0.17 % | - | - | - | - | - |
| *Ruminiclostridium 9* | 58 % | 0.08 % | 0.10 % | - | + | - | - | - |
| *[Clostridium] innocuum* group | 58 % | 0.04 % | 0.05 % | - | + | - | - | - |
| *Ruminiclostridium 5* | 58 % | 0.03 % | 0.04 % | - | + | + | + | - |
| | | | | - | + | + | - | - |
| **Fusobacteria** | **67 %** | **3.50 %** | **5.67 %** | - | + | - | - | - |
| *Fusobacterium* | 67 % | 3.50 % | 5.66 % | - | + | + | - | - |
| **Proteobacteria** | **100 %** | **36.78 %** | **27.90 %** | | | | | |
| *Escherichia-Shigella* | 92 % | 35.19 % | 28.86 % | - | + | - | + | + |
| *Klebsiella* | 58 % | 1.54 % | 3.24 % | | | | | |

***SUPPLEMENTARY TABLE 3: Genus-level bacterial taxa observed in 7 d old foal rectum in >50% of animals.*** *Prevalences, mean relative abundances and standard deviations in 7 d old foals (n=14), and sharing of taxa with the other samples (NB = newborn, FE = mare feces, VA = mare vagina, MO = mare mouth).*

| *Taxon* | *Prevalence* | *Mean* | *SD* | *NB* | *24h* | *FE* | *VA* | *MO* |
|---|---|---|---|---|---|---|---|---|
| **Actinobacteria** | **100 %** | **0.37 %** | **0.31 %** | | | | | |
| *Eggerthella* | 79 % | 0.11 % | 0.13 % | - | - | - | - | - |
| **Bacteroidetes** | **100 %** | **27.86 %** | **9.46 %** | | | | | |
| *Bacteroides* | 100 % | 17.96 % | 9.52 % | + | + | - | + | - |
| *Parabacteroides* | 100 % | 4.43 % | 5.06 % | - | - | - | - | - |
| *Alistipes* | 100 % | 2.51 % | 3.30 % | - | - | - | - | - |
| *Butyricimonas* | 64 % | 0.70 % | 1.45 % | - | - | - | - | - |
| *Odoribacter* | 57 % | 1.08 % | 1.56 % | - | - | - | - | - |
| **Firmicutes** | **100 %** | **55.03 %** | **12.88 %** | | | | | |
| *Tyzzerella 4* | 100 % | 6.16 % | 5.02 % | - | + | - | - | - |
| *Streptococcus* | 100 % | 5.96 % | 3.76 % | + | + | + | + | + |
| *Lactobacillus* | 100 % | 5.46 % | 5.21 % | + | - | + | + | - |
| *Faecalitalea* | 100 % | 3.90 % | 3.36 % | - | + | - | - | - |
| *[Ruminococcus] torques* group | 100 % | 2.55 % | 1.69 % | - | - | - | - | - |
| *Blautia* | 100 % | 2.30 % | 5.71 % | - | - | + | - | - |
| *Flavonifractor* | 100 % | 1.66 % | 1.58 % | - | - | - | - | - |
| *[Eubacterium] coprostanoligenes* group | 100 % | 1.41 % | 0.83 % | + | - | + | - | - |
| *Ruminiclostridium 9* | 100 % | 1.34 % | 0.99 % | - | + | + | - | - |
| *Lachnoclostridium* | 100 % | 1.25 % | 1.12 % | - | - | - | - | - |
| *Erysipelatoclostridium* | 100 % | 1.15 % | 0.75 % | - | + | + | - | - |
| unclassified *Ruminococcaceae* | 100 % | 1.07 % | 1.06 % | - | + | + | + | - |
| *Butyricicoccus* | 100 % | 1.04 % | 0.75 % | - | + | - | - | - |
| *Oscillibacter* | 100 % | 0.67 % | 0.88 % | - | - | + | - | - |
| *Ruminococcaceae UCG-004* | 100 % | 0.56 % | 0.27 % | - | - | + | + | - |
| uncultured *Ruminococcaceae* | 100 % | 0.25 % | 0.58 % | - | - | + | - | - |
| *UBA1819* | 100 % | 0.23 % | 0.16 % | - | - | - | - | - |
| *Ruminiclostridium 5* | 100 % | 0.20 % | 0.20 % | - | + | + | - | - |
| *Fournierella* | 93 % | 4.57 % | 4.05 % | - | + | - | - | - |
| unclassified *Lachnospiraceae* | 93 % | 3.71 % | 3.19 % | + | + | + | + | - |
| *Sellimonas* | 93 % | 1.34 % | 1.45 % | - | - | - | - | - |
| *Tyzzerella* | 93 % | 0.96 % | 0.69 % | - | - | - | - | - |
| *Negativibacillus* | 93 % | 0.54 % | 0.75 % | - | - | - | - | - |
| *Eubacterium* | 93 % | 0.18 % | 0.20 % | - | - | + | - | - |
| *Candidatus Soleaferrea* | 93 % | 0.05 % | 0.04 % | - | - | + | - | - |
| *Phascolarctobacterium* | 86 % | 0.50 % | 0.36 % | + | - | + | - | - |
| *Clostridium sensu stricto 1* | 86 % | 0.27 % | 0.43 % | + | + | + | + | - |
| *Anaerotruncus* | 86 % | 0.21 % | 0.32 % | - | - | - | - | - |
| *Family XIII AD3011* group | 86 % | 0.07 % | 0.06 % | - | - | + | - | - |
| *[Ruminococcus] gnavus* group | 79 % | 0.44 % | 0.57 % | - | + | - | - | - |
| *[Clostridium] innocuum* group | 79 % | 0.28 % | 0.87 % | - | + | - | - | - |
| *[Eubacterium] nodatum* group | 79 % | 0.21 % | 0.22 % | - | - | + | - | - |
| *Pseudoflavonifractor* | 71 % | 0.27 % | 0.34 % | - | - | - | - | - |
| *Eisenbergiella* | 71 % | 0.20 % | 0.33 % | - | - | - | - | - |
| *Intestinimonas* | 71 % | 0.06 % | 0.07 % | - | - | - | - | - |
| *Dielma* | 71 % | 0.04 % | 0.05 % | - | - | - | - | - |
| *[Eubacterium] fissicatena* group | 71 % | 0.03 % | 0.03 % | - | - | - | - | - |
| *Ruminococcaceae UCG-014* | 64 % | 0.61 % | 0.88 % | - | - | + | + | - |
| *Veillonella* | 57 % | 0.26 % | 0.46 % | - | - | - | - | - |
| *Terrisporobacter* | 57 % | 0.21 % | 0.32 % | - | + | - | - | - |
| *Holdemania* | 57 % | 0.03 % | 0.04 % | - | - | - | - | - |
| **Fusobacteria** | **100 %** | **9.60 %** | **8.77 %** | | | | | |

| | | | | | | | | |
|---|---|---|---|---|---|---|---|---|
| *Fusobacterium* | 100 % | 9.60 % | 8.77 % | - | + | - | + | + |
| **Proteobacteria** | **100 %** | **2.62 %** | **2.15 %** | | | | | |
| *Sutterella* | 93 % | 0.63 % | 0.84 % | - | - | + | - | - |
| *Bilophila* | 79 % | 0.29 % | 0.27 % | - | - | - | - | - |
| *Escherichia-Shigella* | 71 % | 0.81 % | 1.21 % | - | + | - | - | - |
| *Desulfovibrio* | 57 % | 0.41 % | 0.44 % | - | - | + | - | - |
| **Verrucomicrobia** | 71 % | 4.36 % | 9.45 % | | | | | |
| *Akkermansia* | 71 % | 4.36 % | 9.45 % | - | - | + | - | - |

#### *Core bacterial taxa in mares*

Most prevalent bacterial taxa in mare feces, vaginal vestibulum and mouth (present in >50% of animals) are shown in Supplementary Tables 4 – 6.

***SUPPLEMENTARY TABLE 4: Genus-level bacterial taxa observed in mare feces in >50% of animals.*** *Prevalences, mean relative abundances and standard deviations in mare feces (n=14), and sharing of taxa with the other samples (NB = newborn, VA = mare vagina, MO = mare mouth).*

| *Taxon* | *Prevalence* | *Mean* | *SD* | *NB* | *24h* | *7d* | *VA* | *MO* |
|---|---|---|---|---|---|---|---|---|
| **Actinobacteria** | **100 %** | **0.69 %** | **0.27 %** | | | | | |
| uncultured *Eggerthellaceae* | 100 % | 0.37 % | 0.16 % | - | - | - | - | - |
| uncultured *Coriobacteriales Incertae Sedis* | 100 % | 0.05 % | 0.04 % | - | - | - | - | - |
| unclassified *Eggerthellaceae* | 93 % | 0.05 % | 0.04 % | - | - | - | - | - |
| uncultured *Coriobacteriales* | 86 % | 0.07 % | 0.05 % | - | - | - | + | - |
| *Phoenicibacter* | 71 % | 0.11 % | 0.10 % | - | - | - | - | - |
| **Armatimonadetes** | **86 %** | **0.05 %** | **0.03 %** | | | | | |
| uncultured *Armatimonadetes* | 86 % | 0.04 % | 0.03 % | - | - | - | - | - |
| **Bacteroidetes** | **100 %** | **24.26 %** | **3.85 %** | | | | | |
| *Rikenellaceae RC9* gut group | 100 % | 7.12 % | 3.25 % | + | - | - | + | - |
| uncultured *Bacteroidales p-251-o5* | 100 % | 3.61 % | 2.38 % | - | - | - | - | - |
| uncultured *Bacteroidales F082* | 100 % | 3.51 % | 2.10 % | - | - | - | - | - |
| uncultured *Bacteroidales RF16* group | 100 % | 1.85 % | 0.64 % | - | - | - | + | - |
| uncultured *Bacteroidales UCG-001* | 100 % | 1.18 % | 0.90 % | - | - | - | - | - |
| *Prevotellaceae UCG-001* | 100 % | 0.97 % | 0.49 % | - | - | - | - | - |
| *Prevotella 1* | 100 % | 0.95 % | 0.50 % | - | - | - | - | - |
| *Prevotellaceae UCG-003* | 100 % | 0.75 % | 0.38 % | - | - | - | - | - |
| *Prevotellaceae UCG-004* | 100 % | 0.44 % | 0.23 % | - | - | - | - | - |
| *hoa5-07d05* gut group | 100 % | 0.41 % | 0.32 % | - | - | - | - | - |
| unclassified *Bacteroidales* | 100 % | 0.37 % | 0.49 % | - | - | - | - | - |
| uncultured *Muribaculaceae* | 100 % | 0.12 % | 0.06 % | - | - | - | - | - |
| uncultured *Paludibacteraceae* | 93 % | 0.74 % | 0.95 % | - | - | - | - | - |
| uncultured *Prevotellaceae* | 93 % | 0.53 % | 1.37 % | - | - | - | - | - |
| uncultured *Bacteroidales BS11* gut group | 93 % | 0.34 % | 0.24 % | - | - | - | - | - |
| uncultured *Marinifilaceae* | 93 % | 0.29 % | 0.20 % | - | - | - | - | - |
| unclassified *Prevotellaceae* | 93 % | 0.21 % | 0.16 % | - | - | - | - | - |
| *Alloprevotella* | 93 % | 0.17 % | 0.11 % | - | - | - | + | - |
| uncultured *Bacteroidales* | 93 % | 0.13 % | 0.23 % | - | - | - | - | - |
| *dgA-11* gut group | 86 % | 0.08 % | 0.06 % | - | - | - | - | - |
| unclassified *Rikenellaceae* | 86 % | 0.05 % | 0.07 % | - | - | - | - | - |
| *Prevotellaceae Ga6A1* group | 64 % | 0.04 % | 0.05 % | - | - | - | - | - |
| *Bacteroidetes bacterium GWF2_40_13* | 57 % | 0.06 % | 0.07 % | - | - | - | - | - |
| uncultured *M2PB4-65 termite* group | 57 % | 0.03 % | 0.05 % | - | - | - | - | - |
| **Cyanobacteria** | **93 %** | **0.30 %** | **0.17 %** | | | | | |
| uncultured *Gastranaerophilales* | 93 % | 0.17 % | 0.10 % | - | - | - | - | - |
| unclassified *Gastranaerophilales* | 93 % | 0.06 % | 0.06 % | - | - | - | - | - |
| rumen bacterium *YS2* | 79 % | 0.04 % | 0.03 % | - | - | - | - | - |
| **Elusimicrobia** | **79 %** | **0.02 %** | **0.03 %** | | | | | |
| *Elusimicrobium* | 71 % | 0.02 % | 0.02 % | - | - | - | - | - |
| **Epsilonbacteraeota** | **64 %** | **0.01 %** | **0.01 %** | | | | | |
| *Campylobacter* | 64 % | 0.01 % | 0.01 % | - | - | - | + | - |
| **Euryarchaeota** | **100 %** | **0.24 %** | **0.22 %** | | | | | |
| *Methanocorpusculum* | 86 % | 0.19 % | 0.21 % | - | - | - | - | - |

| | | | | | | | | |
|---|---|---|---|---|---|---|---|---|
| *Methanobrevibacter* | 79 % | 0.05 % | 0.05 % | - | - | - | - | - |
| **Fibrobacteres** | **100 %** | **2.32 %** | **1.58 %** | | | | | |
| *Fibrobacter* | 100 % | 2.32 % | 1.58 % | - | - | - | - | - |
| **Firmicutes** | **100 %** | **61.38 %** | **3.97 %** | | | | | |
| unclassified *Lachnospiraceae* | 100 % | 8.47 % | 2.77 % | + | + | + | + | - |
| *Ruminococcaceae NK4A214* group | 100 % | 4.46 % | 1.66 % | - | - | - | - | - |
| *Ruminococcus 1* | 100 % | 4.16 % | 1.70 % | - | - | - | - | - |
| *Ruminococcaceae UCG-010* | 100 % | 3.95 % | 1.99 % | + | - | - | + | - |
| *Christensenellaceae R-7* group | 100 % | 3.48 % | 1.59 % | - | - | - | + | - |
| *Lachnospiraceae XPB1014* group | 100 % | 3.16 % | 1.40 % | - | - | - | - | - |
| *Lachnospiraceae AC2044* group | 100 % | 2.64 % | 1.00 % | - | - | - | - | - |
| *Ruminococcaceae UCG-005* | 100 % | 2.62 % | 1.11 % | + | - | - | - | - |
| *[Eubacterium] coprostanoligenes* group | 100 % | 2.36 % | 1.16 % | + | - | + | - | - |
| *Marvinbryantia* | 100 % | 1.99 % | 1.00 % | - | - | - | - | - |
| *Ruminococcaceae UCG-002* | 100 % | 1.72 % | 1.10 % | - | - | - | + | - |
| *Lachnospiraceae UCG-009* | 100 % | 1.31 % | 0.62 % | - | - | - | - | - |
| *Ruminiclostridium 9* | 100 % | 1.14 % | 1.09 % | - | + | + | - | - |
| *[Eubacterium] hallii* group | 100 % | 1.13 % | 0.63 % | - | - | - | - | - |
| *Blautia* | 100 % | 1.09 % | 0.63 % | - | - | + | - | - |
| *Family XIII AD3011* group | 100 % | 1.07 % | 0.37 % | - | - | + | - | - |
| *Anaerovorax* | 100 % | 0.92 % | 0.64 % | + | - | - | - | - |
| *Phascolarctobacterium* | 100 % | 0.89 % | 0.28 % | + | - | + | - | - |
| unclassified *Ruminococcaceae* | 100 % | 0.86 % | 0.43 % | - | + | + | + | - |
| *Saccharofermentans* | 100 % | 0.74 % | 0.32 % | - | - | - | - | - |
| uncultured *Lachnospiraceae* | 100 % | 0.57 % | 0.34 % | - | - | - | - | - |
| uncultured *Ruminococcaceae* | 100 % | 0.57 % | 0.25 % | - | - | + | - | - |
| *Erysipelotrichaceae UCG-004* | 100 % | 0.55 % | 0.38 % | - | - | - | - | - |
| *Lachnospiraceae NK4A136* group | 100 % | 0.53 % | 0.23 % | - | - | - | - | - |
| *Candidatus Soleaferrea* | 100 % | 0.52 % | 0.45 % | - | - | + | - | - |
| *Papillibacter* | 100 % | 0.50 % | 0.47 % | - | - | - | - | - |
| *Ruminococcaceae UCG-014* | 100 % | 0.50 % | 0.23 % | - | - | + | + | - |
| *Oribacterium* | 100 % | 0.41 % | 0.21 % | - | - | - | - | - |
| *Agathobacter* | 100 % | 0.37 % | 0.21 % | - | - | - | - | - |
| *[Eubacterium] ruminantium* group | 100 % | 0.35 % | 0.18 % | - | - | - | - | - |
| *Catenisphaera* | 100 % | 0.35 % | 0.28 % | - | - | - | - | - |
| *Ruminococcaceae UCG-004* | 100 % | 0.33 % | 0.16 % | - | - | + | + | - |
| *Ruminococcaceae UCG-013* | 100 % | 0.31 % | 0.19 % | - | - | - | - | - |
| uncultured *Clostridiales vadinBB60* group | 100 % | 0.31 % | 0.16 % | - | - | - | - | - |
| *Lactobacillus* | 100 % | 0.30 % | 0.20 % | + | - | + | + | - |
| unclassified *Family XIII* | 100 % | 0.28 % | 0.20 % | - | - | - | - | - |
| *Sarcina* | 100 % | 0.27 % | 0.23 % | - | - | - | - | - |
| *Family XIII UCG-001* | 100 % | 0.27 % | 0.09 % | - | - | - | - | - |
| *Pseudobutyrivibrio* | 100 % | 0.27 % | 0.17 % | - | - | - | - | - |
| *Anaerovibrio* | 100 % | 0.20 % | 0.15 % | - | - | - | - | - |
| *Mogibacterium* | 100 % | 0.19 % | 0.14 % | - | - | - | + | - |
| *Defluviitaleaceae UCG-011* | 100 % | 0.18 % | 0.07 % | - | - | - | - | - |
| uncultured *Peptococcaceae* | 100 % | 0.18 % | 0.07 % | - | - | - | - | - |
| uncultured *Family XIII* | 100 % | 0.16 % | 0.08 % | - | - | - | - | - |
| *Lachnoclostridium 10* | 100 % | 0.16 % | 0.08 % | - | - | - | - | - |
| *Ruminococcaceae UCG-007* | 100 % | 0.15 % | 0.08 % | - | - | - | - | - |
| *Acetitomaculum* | 100 % | 0.15 % | 0.14 % | - | - | - | - | - |
| uncultured *Erysipelotrichaceae* | 100 % | 0.13 % | 0.05 % | - | - | - | - | - |
| *Quinella* | 100 % | 0.12 % | 0.09 % | - | - | - | - | - |
| *Lachnospiraceae UCG-008* | 100 % | 0.12 % | 0.08 % | - | - | - | - | - |
| *Lachnospiraceae NK4B4* group | 100 % | 0.10 % | 0.04 % | - | - | - | - | - |
| unclassified *Erysipelotrichaceae* | 100 % | 0.10 % | 0.08 % | - | - | - | - | - |
| uncultured *Christensenellaceae* | 100 % | 0.06 % | 0.04 % | - | - | - | - | - |
| *Eubacterium* | 93 % | 0.40 % | 0.35 % | - | - | + | - | - |
| unclassified *Clostridiales* | 93 % | 0.15 % | 0.22 % | - | - | - | - | - |

| | | | | | | | | |
|---|---|---|---|---|---|---|---|---|
| *Oscillibacter* | 93 % | 0.13 % | 0.13 % | - | - | + | - | - |
| *Oscillospira* | 93 % | 0.09 % | 0.07 % | - | - | - | - | - |
| *Lachnospiraceae FCS020* group | 93 % | 0.09 % | 0.06 % | - | - | - | - | - |
| *[Eubacterium] nodatum* group | 93 % | 0.08 % | 0.06 % | - | - | + | - | - |
| *Erysipelatoclostridium* | 93 % | 0.07 % | 0.06 % | - | + | + | - | - |
| *Shuttleworthia* | 93 % | 0.06 % | 0.04 % | - | - | - | - | - |
| *Clostridium sensu stricto 1* | 86 % | 0.24 % | 0.29 % | + | + | + | + | - |
| *Lachnospiraceae ND3007* group | 86 % | 0.11 % | 0.07 % | - | - | - | - | - |
| *Hydrogenoanaerobacterium* | 86 % | 0.11 % | 0.13 % | - | - | - | - | - |
| uncultured *Veillonellaceae* | 86 % | 0.09 % | 0.15 % | - | - | - | - | - |
| *Coprococcus 2* | 86 % | 0.09 % | 0.07 % | - | - | - | - | - |
| *Lachnospiraceae UCG-006* | 86 % | 0.07 % | 0.04 % | - | - | - | - | - |
| *Clostridiales vadinBB60* group rumen bacterium | 86 % | 0.06 % | 0.06 % | - | - | - | - | - |
| *Ruminococcaceae UCG-009* | 86 % | 0.06 % | 0.05 % | - | - | - | - | - |
| *Coprococcus 1* | 86 % | 0.04 % | 0.04 % | - | - | - | - | - |
| *Faecalibacterium* | 86 % | 0.04 % | 0.02 % | - | - | - | - | - |
| *Pygmaiobacter* | 79 % | 0.07 % | 0.10 % | - | - | - | - | - |
| *Lachnospiraceae UCG-002* | 79 % | 0.06 % | 0.06 % | - | - | - | - | - |
| *Ruminococcaceae V9D2013* group | 79 % | 0.06 % | 0.07 % | - | - | - | - | - |
| unclassified *Clostridiales vadinBB60* group | 79 % | 0.06 % | 0.05 % | - | - | - | - | - |
| *Lachnospiraceae FE2018* group | 79 % | 0.04 % | 0.03 % | - | - | - | - | - |
| *Ruminiclostridium 5* | 79 % | 0.04 % | 0.03 % | - | + | + | - | - |
| *Clostridiales vadinBB60* group metagenome | 79 % | 0.04 % | 0.03 % | - | - | - | - | - |
| *Streptococcus* | 71 % | 0.48 % | 1.17 % | + | + | + | + | + |
| *FD2005* | 71 % | 0.06 % | 0.05 % | - | - | - | - | - |
| *Ruminiclostridium 1* | 71 % | 0.05 % | 0.05 % | - | - | - | - | - |
| *Subdoligranulum* | 71 % | 0.04 % | 0.04 % | - | - | - | - | - |
| *Johnsonella* | 71 % | 0.03 % | 0.03 % | - | - | - | - | - |
| *Ruminococcus 2* | 64 % | 0.08 % | 0.16 % | - | - | - | - | - |
| *Cellulosilyticum* | 64 % | 0.06 % | 0.07 % | - | - | - | - | - |
| *[Eubacterium] saphenum* group | 64 % | 0.03 % | 0.05 % | - | - | - | - | - |
| *Moryella* | 64 % | 0.03 % | 0.02 % | - | - | - | - | - |
| *Erysipelotrichaceae UCG-003* | 64 % | 0.02 % | 0.02 % | - | - | - | - | - |
| **Kiritimatiellaeota** | **100 %** | **5.00 %** | **1.83 %** | | | | | |
| uncultured *WCHB1-41* | 100 % | 3.63 % | 1.52 % | + | - | - | - | - |
| unclassified *WCHB1-41* | 100 % | 0.75 % | 0.28 % | - | - | - | - | - |
| uncultured *WCHB1-41* rumen bacterium | 100 % | 0.62 % | 0.41 % | - | - | - | - | - |
| **Lentisphaerae** | **100 %** | **0.32 %** | **0.39 %** | | | | | |
| uncultured *Victivallales vadinBE97* | 93 % | 0.18 % | 0.31 % | - | - | - | - | - |
| *horsej-a03* | 86 % | 0.06 % | 0.08 % | - | - | - | - | - |
| uncultured *Victivallaceae* | 86 % | 0.04 % | 0.03 % | - | - | - | - | - |
| *Z20* | 64 % | 0.03 % | 0.04 % | - | - | - | - | - |
| **Patescibacteria** | **100 %** | **0.12 %** | **0.10 %** | | | | | |
| *Candidatus Saccharimonas* | 100 % | 0.12 % | 0.10 % | - | - | - | - | - |
| **Planctomycetes** | **100 %** | **0.06 %** | **0.03 %** | | | | | |
| *p-1088-a5* gut group | 100 % | 0.05 % | 0.03 % | - | - | - | - | - |
| **Proteobacteria** | **100 %** | **0.66 %** | **0.26 %** | | | | | |
| uncultured *Rhodospirillales* rumen bacterium | 100 % | 0.11 % | 0.07 % | - | - | - | - | - |
| *Sutterella* | 93 % | 0.03 % | 0.02 % | - | - | + | - | - |
| uncultured *Rhodospirillales bacterium* | 86 % | 0.16 % | 0.19 % | - | - | - | - | - |
| *Desulfovibrio* | 86 % | 0.11 % | 0.11 % | - | - | + | - | - |
| *Mailhella* | 79 % | 0.06 % | 0.06 % | - | - | - | - | - |
| *Rhodospirillales gut* metagenome | 64 % | 0.02 % | 0.02 % | - | - | - | - | - |
| **Spirochaetes** | **100 %** | **3.37 %** | **2.67 %** | | | | | |

| | | | | | | | | |
|---|---|---|---|---|---|---|---|---|
| *Treponema 2* | 100 % | 3.07 % | 2.67 % | - | - | - | - | - |
| uncultured *MVP-15* | 71 % | 0.07 % | 0.16 % | - | - | - | - | - |
| *Sediminispirochaeta* | 71 % | 0.02 % | 0.03 % | - | - | - | - | - |
| **Synergistetes** | **93 %** | **0.24 %** | **0.33 %** | | | | | |
| uncultured *Synergistaceae* | 79 % | 0.09 % | 0.17 % | - | - | - | - | - |
| *Cloacibacillus* | 71 % | 0.09 % | 0.14 % | - | - | - | - | - |
| *Pyramidobacter* | 64 % | 0.03 % | 0.04 % | - | - | - | - | - |
| **Tenericutes** | **100 %** | **0.20 %** | **0.13 %** | | | | | |
| *Anaeroplasma* | 93 % | 0.07 % | 0.06 % | - | - | - | - | - |
| unclassified *Mollicutes RF39* | 64 % | 0.05 % | 0.05 % | - | - | - | - | - |
| uncultured *Mollicutes F39* | 57 % | 0.03 % | 0.03 % | - | - | - | - | - |
| **Verrucomicrobia** | **100 %** | **0.66 %** | **0.88 %** | | | | | |
| *Akkermansia* | 100 % | 0.61 % | 0.85 % | - | - | + | - | - |

***SUPPLEMENTARY TABLE 5: Genus-level bacterial taxa observed in mare vagina in >50% of animals.***
*Prevalences, mean relative abundances and standard deviations in mare vaginal vestibulum (n=14), and sharing of taxa with the other samples (NB = newborn, FE = mare feces, MO = mare mouth).*

| *Taxon* | *Prevalence* | *Mean* | *SD* | *NB* | *24h* | *7d* | *FE* | *MO* |
|---|---|---|---|---|---|---|---|---|
| **Actinobacteria** | **100 %** | **24.48 %** | **15.88 %** | - | - | - | - | - |
| *Corynebacterium* | 100 % | 17.60 % | 14.96 % | - | - | - | - | - |
| *Arcanobacterium* | 100 % | 1.87 % | 2.80 % | - | - | - | - | - |
| uncultured *Propionibacteriaceae* | 93 % | 0.14 % | 0.15 % | - | - | - | - | - |
| *Corynebacterium 1* | 86 % | 2.44 % | 5.98 % | + | - | - | - | - |
| unclassified *Corynebacteriaceae* | 71 % | 0.57 % | 1.04 % | - | - | - | - | - |
| *Mobiluncus* | 64 % | 0.36 % | 0.70 % | - | - | - | - | - |
| uncultured *Coriobacteriales* | 64 % | 0.11 % | 0.17 % | - | - | - | + | - |
| **Bacteroidetes** | **100 %** | **18.97 %** | **11.44 %** | | | | | |
| *Porphyromonas* | 100 % | 14.25 % | 11.08 % | - | - | - | - | + |
| *Bacteroides* | 64 % | 0.16 % | 0.48 % | + | + | + | - | - |
| *Alloprevotella* | 64 % | 0.15 % | 0.43 % | - | - | - | + | - |
| *Rikenellaceae RC9* gut group | 57 % | 0.93 % | 2.58 % | + | - | - | + | - |
| uncultured *Bacteroidales RF16* group | 57 % | 0.27 % | 0.70 % | - | - | - | + | - |
| **Epsilonbacteraeota** | **100 %** | **8.45 %** | **4.89 %** | | | | | |
| *Campylobacter* | 100 % | 8.45 % | 4.89 % | - | - | - | + | - |
| **Firmicutes** | **100 %** | **36.73 %** | **11.88 %** | | | | | |
| *Helcococcus* | 100 % | 8.97 % | 8.77 % | - | - | - | - | - |
| *Streptococcus* | 100 % | 6.18 % | 9.28 % | + | + | + | + | + |
| *Peptoniphilus* | 100 % | 3.96 % | 2.99 % | - | - | - | - | - |
| *Globicatella* | 100 % | 3.16 % | 2.46 % | - | - | - | - | - |
| unclassified *Ruminococcaceae* | 93 % | 0.35 % | 0.74 % | - | + | + | + | - |
| *Peptococcus* | 93 % | 0.33 % | 0.43 % | - | - | - | - | - |
| *Anaerococcus* | 86 % | 2.97 % | 6.10 % | - | - | - | - | - |
| *Peptostreptococcus* | 86 % | 1.87 % | 1.96 % | - | - | - | - | - |
| unclassified *Peptostreptococcaceae* | 86 % | 0.76 % | 1.53 % | - | - | - | - | - |
| *Murdochiella* | 79 % | 1.39 % | 1.42 % | - | - | - | - | - |
| unclassified *Lachnospiraceae* | 71 % | 0.27 % | 0.38 % | + | + | + | + | - |
| *Christensenellaceae R-7* group | 71 % | 0.27 % | 0.51 % | - | - | - | + | - |
| *Clostridium sensu stricto 1* | 71 % | 0.06 % | 0.07 % | + | + | + | + | - |
| *Ruminococcaceae UCG-002* | 64 % | 0.35 % | 0.81 % | - | - | - | + | - |
| *Staphylococcus* | 64 % | 0.31 % | 0.69 % | + | - | - | - | - |
| *Ignavigranum* | 64 % | 0.18 % | 0.33 % | - | - | - | - | - |
| *Bacillus* | 64 % | 0.17 % | 0.38 % | + | - | - | - | - |
| *Mogibacterium* | 64 % | 0.10 % | 0.21 % | - | - | - | + | - |
| *Ruminococcaceae UCG-004* | 64 % | 0.06 % | 0.12 % | - | - | + | + | - |
| *Ruminococcaceae UCG-010* | 57 % | 0.58 % | 1.64 % | + | - | - | + | - |
| *Ruminococcaceae UCG-014* | 57 % | 0.12 % | 0.21 % | - | - | + | + | - |
| *Lactobacillus* | 57 % | 0.02 % | 0.03 % | + | - | + | + | - |
| **Fusobacteria** | **100 %** | **5.64 %** | **5.68 %** | | | | | |
| *Oceanivirga* | 86 % | 4.41 % | 5.90 % | - | - | - | - | - |
| *Fusobacterium* | 71 % | 1.17 % | 2.73 % | - | + | + | - | + |
| **Proteobacteria** | **93 %** | **3.61 %** | **8.91 %** | | | | | |
| *Sphingomonas* | 57 % | 0.42 % | 0.94 % | + | - | - | - | - |

***SUPPLEMENTARY TABLE 6: Genus-level bacterial taxa observed in mare mouth in >50% of animals.*** *Prevalences, mean relative abundances and standard deviations in mare vestibulum oris (n=14), and sharing of taxa with the other samples (NB = newborn, FE = mare feces, VA = mare vagina).*

| *Taxon* | *Prevalence* | *Mean* | *SD* | *NB* | *24h* | *7d* | *FE* | *VA* |
|---|---|---|---|---|---|---|---|---|
| **Bacteroidetes** | **100 %** | **1.03 %** | **1.86 %** | | | | | |
| *Porphyromonas* | 86 % | 0.43 % | 0.90 % | - | - | - | - | + |
| *Bergeyella* | 79 % | 0.13 % | 0.17 % | - | - | - | - | - |
| **Firmicutes** | **100 %** | **81.28 %** | **15.12 %** | | | | | |
| *Gemella* | 100 % | 74.13 % | 16.23 % | - | - | - | - | - |
| *Streptococcus* | 100 % | 4.71 % | 5.11 % | + | + | + | + | + |
| *Enterococcus* | 100 % | 1.66 % | 3.33 % | - | + | - | - | - |
| **Fusobacteria** | **93 %** | **0.16 %** | **0.16 %** | | | | | |
| *Leptotrichia* | 79 % | 0.12 % | 0.13 % | - | - | - | - | - |
| *Fusobacterium* | 79 % | 0.04 % | 0.05 % | - | + | + | - | + |
| **Proteobacteria** | **100 %** | **15.89 %** | **13.09 %** | | | | | |
| unclassified *Pasteurellaceae* | 100 % | 5.74 % | 8.68 % | - | - | - | - | - |
| *Moraxella* | 100 % | 2.81 % | 5.91 % | - | - | - | - | - |
| *Actinobacillus* | 93 % | 1.99 % | 4.57 % | - | - | - | - | - |
| *Neisseria* | 93 % | 0.30 % | 0.45 % | - | - | - | - | - |
| *Alysiella* | 86 % | 3.88 % | 8.03 % | - | - | - | - | - |
| *Lautropia* | 86 % | 0.17 % | 0.18 % | - | - | - | - | - |

#### *Comparison of foal and mare microbiotas*

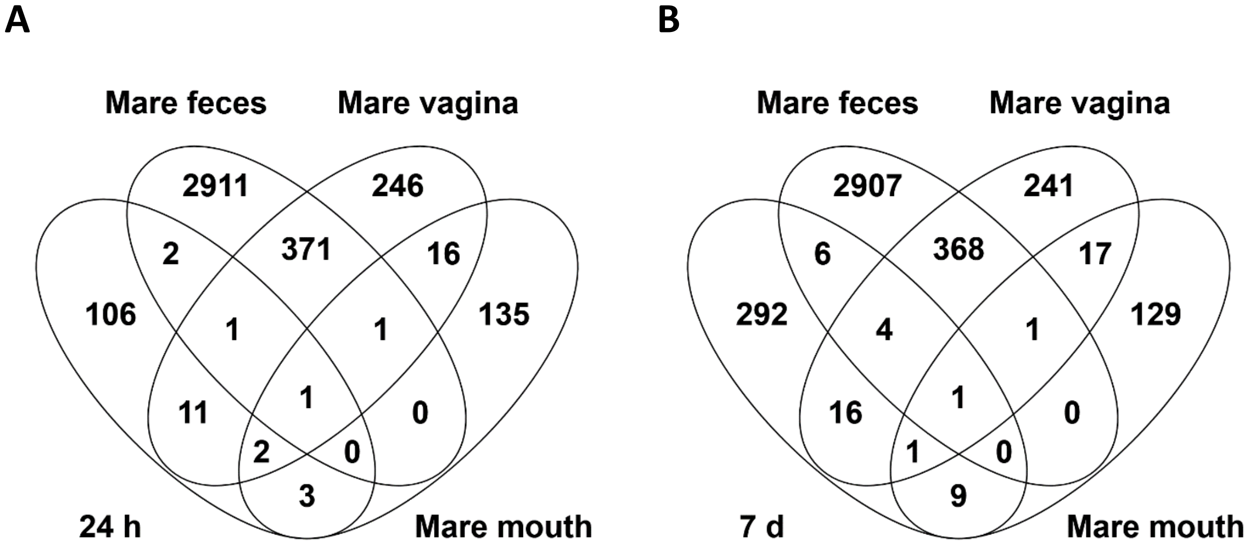

***SUPPLEMENTARY FIGURE 3.*** *ASVs shared between rectal microbiota samples of 24 h foals (A) and 7 d foals (B) and various mare microbiota samples. All ASVs detected in at least 2 animals per sample group are included.*

The 7-day rectal microbiota clustered closer to the adult fecal microbiota at the family level than at the ASV level (Supplementary Fig. 4).

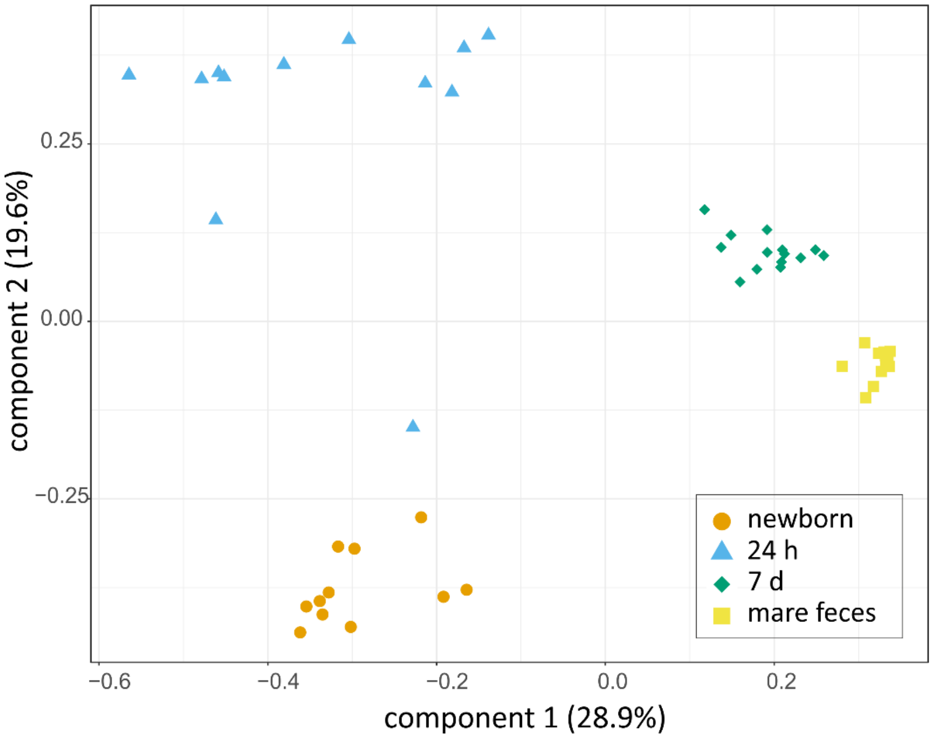

***SUPPLEMENTARY FIGURE 4.*** *PCoA on Bray-Curtis distances of rectal microbiota in the newborn, 24 h old and 7 d old foals and adult mares at the family level. Colors and shapes indicate the sample types.*

### Supplementary methods

#### *Quantitative PCR*

A universal probe and primer set targeting the bacterial 16S rRNA gene was used for the quantification of 16S rRNA gene copy numbers in the DNA extracted from the foal rectal samples (newborn, 24 h and 7 d) and negative controls (no-template, instrument and field controls) [forward primer: 5′-TCCTACGGGAGGCAGCAGT-3′; reverse primer: 5′-GGACTACCAGGGTATCTAATCCTGTT-3′; probe: (6-FAM)-5′-CGTATTACCGCGGCTGCTGGCAC-3′-(BHQ1)][1]. To prepare a standard curve for absolute quantification, the complete 16S rRNA gene from *Lactobacillus amylovorus* (strain GRL1112) was amplified using primers 5′-AGAGTTTGATCCTGGCTCAG-3′ and 5′-ACGGCTACCTTGTTACGACTT-3′[2]. A standard series consisting of seven ten-fold dilutions from $1x10^2$ to $1x10^8$ copies was used.

The qPCR was performed with an Mx3005P instrument (Agilent Technologies, Santa Clara, CA, USA) using DNA free 96 well plates (Eppendorf, Germany). The PCR reaction (25 µl total volume) containing 200 nM of probe and 300 nM of each primer was performed utilising the ROX-containing 5× HOT FIREPol Probe qPCR Mix Plus (Solis BioDyne, Estonia). All samples were amplified in triplicate by the following thermal cycling conditions: 95°C for 15 min (polymerase activation and initial template denaturation), followed by 40 cycles of 95°C for 15 s (denaturation) and 60°C for 60 s (annealing/elongation).

To fit all samples within the dynamic range of qPCR, 2.5 µl of control and newborn DNA extract and 0.05 µl of 1 d and 7 d DNA extracts were used per reaction. Fecal and especially the meconium samples contain high concentrations of PCR inhibitors, which are not completely removed by DNA extraction kits. Smaller PCR inhibitor concentrations in the more diluted samples may allow more efficient PCR amplification, resulting in values proportionally up to ~50% higher in the 24 h and 7 d samples, based on dilution series tests. The MxPro – Mx3000P software version 4.10 (Agilent Technologies) was used for data analysis.

##### *MiSeq amplicon sequencing of 16S rRNA genes*

The hypervariable regions V3-V4 of the 16S rRNA genes were sequenced using the Illumina MiSeq platform in the DNA core facility of the University of Helsinki, as described previously[3].

Each sample was first amplified in triplicate in 25 µl total volume, using 1× Phusion Hot Start II High-Fidelity PCR Master Mix (Thermo Scientific), 2.5% DMSO (Thermo Scientific), 500 nM of a mixture of 4 forward and 4 reverse universal bacterial primers (Metabion, Supplementary table 7), and 2.5 µl of DNA extracted from each sample, diluted with Nuclease-free Water 1:1 (Ambion™, Thermo Fisher Scientific, USA). DNA and PCR inhibitor free tubes (STARLAB International, Germany) were used. The newborn samples were ran in the same batch as the negative controls and a no-template control was added to every batch. Otherwise every sample type was ran as its own batch, with the added no-template controls.

The thermal cycling conditions included an initial denaturation step at 98 °C for 30 seconds, followed by an optimized amount of cycles dependent on the sample type of denaturing at 98 °C for 10 seconds, annealing at 56 °C for 30 seconds and extension at 72 °C for 20 seconds. The final extension step was at 72 °C for 5 minutes. T100™ Thermal Cycler (Bio-Rad Laboratories) was used. The newborn meconium samples, instrument controls and field controls were amplified with 21 PCR cycles, mare mouth and vaginal samples with 18 cycles, 24 h foal rectal samples with 16 cycles and 7 d old foal rectal samples and mare fecal samples with 14 cycles. ZymoBIOMICS Microbial Community Standard (Zymo Research, U.S.A.) was amplified with 12 cycles.

After the 1$^{st}$ round PCR, the triplicates were combined to a same tube (STARLAB International, Germany), checked on agarose gel and used as templates for the 2$^{nd}$ round PCR.

The 2$^{nd}$ round PCR amplifications were performed using an Illumina forward and reverse primer set (SUPPLEMENTARY TABLE VII), Phusion Hot-Start II polymerase (Finnzymes/Thermo Scientific), High Fidelity buffer and 2.5 % DMSO. The following thermal cycling conditions were applied with an Arktik thermal cycler (Finnzymes/Thermo Scientific): initial denaturation at 98 °C for 30 s, 17 cycles at 98 °C for 10s, 65°C for 30s, 72 °C for 10s, and a final extension at 72 °C for 5 min.

The second round PCR products were pooled in equal amounts, purified with Agencourt AMPure XP magnetic beads (Beckman Coulter) and size selected (500-700 bp) using BluePippin™ (Sage Science, USA). The quantity and quality of the amplicons were assessed with Qubit (Invitrogen, Thermo Scientific) and Bioanalyzer 2100 (Agilent Technologies), respectively. The final 16S rRNA gene amplicons obtained from samples, negative controls and PCR blanks were sequenced on an Illumina MiSeq sequencer using the v2 600 cycle kit paired-end (325 bp + 285 bp).

***SUPPLEMENTARY TABLE 7. Sequencing primers***

| | | |
|---|---|---|
| 1st round | FW 5'-3' (341F) | ACACTCTTTCCCTACACGACGCTCTTCCGATCT*CCTACGGGNGGCWGCAG* |
| | | ACACTCTTTCCCTACACGACGCTCTTCCGATCT**gt***CCTACGGGNGGCWGCAG* |
| | | ACACTCTTTCCCTACACGACGCTCTTCCGATCT**agag***CCTACGGGNGGCWGCAG* |
| | | ACACTCTTTCCCTACACGACGCTCTTCCGATCT**tagtgt***CCTACGGGNGGCWGCAG* |
| | REV 5'-3' (785R) | GTGACTGGAGTTCAGACGTGTGCTCTTCCGATCT*GACTACHVGGGTATCTAATCC* |
| | | GTGACTGGAGTTCAGACGTGTGCTCTTCCGATCT**a***GACTACHVGGGTATCTAATCC* |
| | | GTGACTGGAGTTCAGACGTGTGCTCTTCCGATCT**tct***GACTACHVGGGTATCTAATCC* |
| | | GTGACTGGAGTTCAGACGTGTGCTCTTCCGATCT**ctgagtg***GACTACHVGGGTATCTAATCC* |
| 2nd round | FW 5'-3' | AATGATACGGCGACCACCGAGATCTACACTCTTTCCCTACACGAC |
| | REV 5'-3' | CAAGCAGAAGACGGCATACGAGAT**XXXXXXXX**GTGACTGGAGTTCAGACGTGT |

▶**Sequencing primer** ▶*16S specific primer* ▶**Spacer nucleotide** ▶Illumina adapter ▶Inde**X**

#### *Detailed description of the bioinformatics pipeline*

##### Obtaining the data

The sequencing data for 108 samples was obtained from the sequencing lab in demultiplexed FASTQ format. The compressed .tar.gz-package was uploaded to supercluster Taito of Finnish Center for Scientific Computing (CSC). Package md5sum was validated before further decompression and analyses.

##### Quality check

FastQC 0.11.8 was ran for all the files and the resulting reports were checked manually[4]. FastQC reports were also compiled and assessed with MultiQC 1.7[5].

##### Trimming

All leftover primers and spacers were removed with Cutadapt v.1.10[6].

```
find -name "*R1_001.fastq.gz" -exec cutadapt -g CCTACGGGNGGCWGCAG -o '{}.TRIMMED_CUTADAPT_FW.gz' '{}' ';'
```

```
find -name "*R2_001.fastq.gz" -exec cutadapt -g GACTACHVGGGTATCTAATCC -o '{}.TRIMMED_CUTADAPT_REV.gz' '{}' ';'
```

The trimmed sequences were again checked with FastQC 0.11.8 and MultiQC 1.7[5].

##### Mapping file

A QIIME 2 compatible mapping containing the sample metadata file was created with Google Sheets and validated with Keemei[7 8].

##### QIIME2

Fastq.gz files were first imported to QIIME2 v2018.8 with command `qiime tools import`[7]. The sequences were checked again using `qiime demux summarize`. The total number of raw sequences was 15242693 (mean per sample: 141136) with quality drop in the forward reads at position 302 and in the reverse reads at position 257. DADA2[9] was run using the command `dada2 denoise-paired` with truncating option F302 and R257. DADA2 pipeline plugin in QIIME2 goes through 1000000 reads to estimate the error model, filters and trims the data, dereplicates, learns error rates, infers sample composition, merges paired reads, makes a sequence table and removes chimaeras[7,9]. This results in an amplicon sequence variant (ASV) table. After DADA2 the total number of sequences was 6393231 (mean per sample: 59197). Approximately 42% of raw reads were preserved.

Stats from the denoising and merging steps were checked with `qiime metadata tabulate`. Resulting data was explored with visual summaries created with `qiime feature-table summarize` and `qiime feature-table tabulate-seqs`. The whole dataset contained 13756 ASVs with a total frequency of 6393231. Median and mean frequency per sample were respectively 60272 and 59196. Median and mean frequency per ASV were 27 and 465.

The phylogenetic tree was created using `qiime alignment mafft`[10]. Highly variable positions adding noise were masked with `qiime alignment mask`. FastTree was used to create a phylogenetic tree from the masked alignment: `qiime phylogeny fasttree` and `qiime phylogeny midpoint-root`[11].

Preliminary diversity analysis was created with `qiime diversity core-metrics-phylogenetic` using sampling depth 37149.

The taxonomy was assigned according to the SILVA v132 QIIME release 97%[12]. Representative set of sequences and taxonomy were imported to QIIME2 with `qiime tools import`. Reference reads were first extracted from the sequence directory with `qiime feature-classifier extract-reads`[13]. Then a Naïve Bayes classifier was trained to the curated taxonomy with `qiime feature-classifier fit-classifier-naïve-bayes`[14]. Finally the actual classification was called with `qiime feature-classifier classify-sklearn`[15].

The taxonomy was visualized again with `qiime metadata tabulate` and `qiime taxa barplot`. After this the ASV table was exported from QIIME2 for handling in spreadsheet programs. First the table and taxonomy data was exported in biom-format with `qiime tools export`[16]. The two were combined with biom package: `biom add-metadata`. Finally the ASV table and table with taxonomy were converted to .tsv format with `biom convert`.

##### Data decontamination

We first removed unassigned ASVs, unclassified bacteria (mostly mitochondrial sequences), chloroplasts and the ASVs which were detected less than 10 times in the entire dataset. The data was then filtered to remove ASVs which represented probable contaminants. An ASV was removed if its prevalence in actual samples was ≤ 2× its prevalence in instrument controls *or* its prevalence in field controls, *or* if it's mean relative abundance in actual samples was ≤ 10× its mean abundance in instrument controls *or* its mean abundance in field controls. The filtering was performed for each sample group separately.
